# Two parallel arms of the heterochronic pathway direct coordinated juvenile-to-adult transition through distinct LIN-29 isoforms

**DOI:** 10.1101/783001

**Authors:** Chiara Azzi, Florian Aeschimann, Anca Neagu, Helge Großhans

## Abstract

Robust organismal development relies on temporal coordination of disparate physiological processes. In *Caenorhabditis elegans*, the timely transition from juvenile to adult is controlled by the heterochronic pathway, a regulatory cascade of conserved proteins and small RNAs. The heterochronic pathway culminates in accumulation of the transcription factor LIN-29, which triggers coordinated execution of juvenile-to-adult (J/A) transition events. Here, we reveal that two LIN-29 isoforms fulfill distinct functions during the J/A transition. We show that the functional differences between the isoforms do not stem from differences in their sequences, but from their distinct spatiotemporal expression, and we propose that distinct LIN-29 dose sensitivities of the individual J/A transition events help to ensure their temporal ordering. We demonstrate that unique *lin-29* isoform expression patterns are generated by the RNA-binding protein LIN-41 for *lin-29a*, and the transcription factor HBL-1 for *lin-29b*. By regulating both HBL-1 and LIN-41, the RNA-binding protein LIN-28 coordinates LIN-29 isoform activity. Thus, our findings reveal that a coordinated transition from juvenile to adult involves branching of a linear pathway to achieve timely control of multiple events.

## Introduction

Temporal coordination of diverse events is a hallmark of organismal development. This is illustrated by the juvenile-to-adult (J/A) transition of animals, in mammals also known as puberty. J/A transition involves coordinated morphological changes of sexual organs as well as various other tissues and organs, including skin (Lee and Houk, 2007). The molecular mechanisms that control the onset of J/A transition in humans are poorly understood, but have been well studied in the nematode *Caenorhabditis elegans* (Faunes and Larraín, 2016). In *C. elegans*, J/A transition is controlled by a cascade of regulators termed the heterochronic pathway, which coordinates somatic cell fate programs (Ambros and Horvitz, 1984). Orthologues of heterochronic genes have also been implicated in timing the onset of puberty in mammals including humans (Abreu et al., 2013; Corre et al., 2016; Ong et al., 2009; Perry et al., 2009; Sulem et al., 2009; Zhu et al., 2010) (reviewed in Faunes and Larraín, 2016; Moss and Romer-Seibert, 2014), indicating an evolutionary conservation of the molecular principles of temporal coordination of J/A transition.

The *C. elegans* J/A transition has been particularly well studied in the epidermis, where it encompasses four events related to cell fates and molting (Figure 1A; (Ambros, 1989)). First, skin progenitor cells called seam cells undergo asymmetric (self-renewal) divisions during larval stages, but cease to do so in adult animals. The last seam cell division takes place during the transition from the third (L3) to the last (L4) larval stage. Second, during the mid-L4 stage, seam cells fuse into a syncytium, a state considered terminally differentiated (Sulston and Horvitz, 1977). Third, animals generate an adult cuticle, characterized by the presence of adult-specific collagens (Cox and Hirsh, 1985), and a microscopically visible structure known as adult alae (Singh and Sulston, 1978). Fourth, following shedding of the L4 cuticle, animals stop molting, the process of cuticle synthesis and shedding that happens at the end of each larval stage.

**Figure 1:**
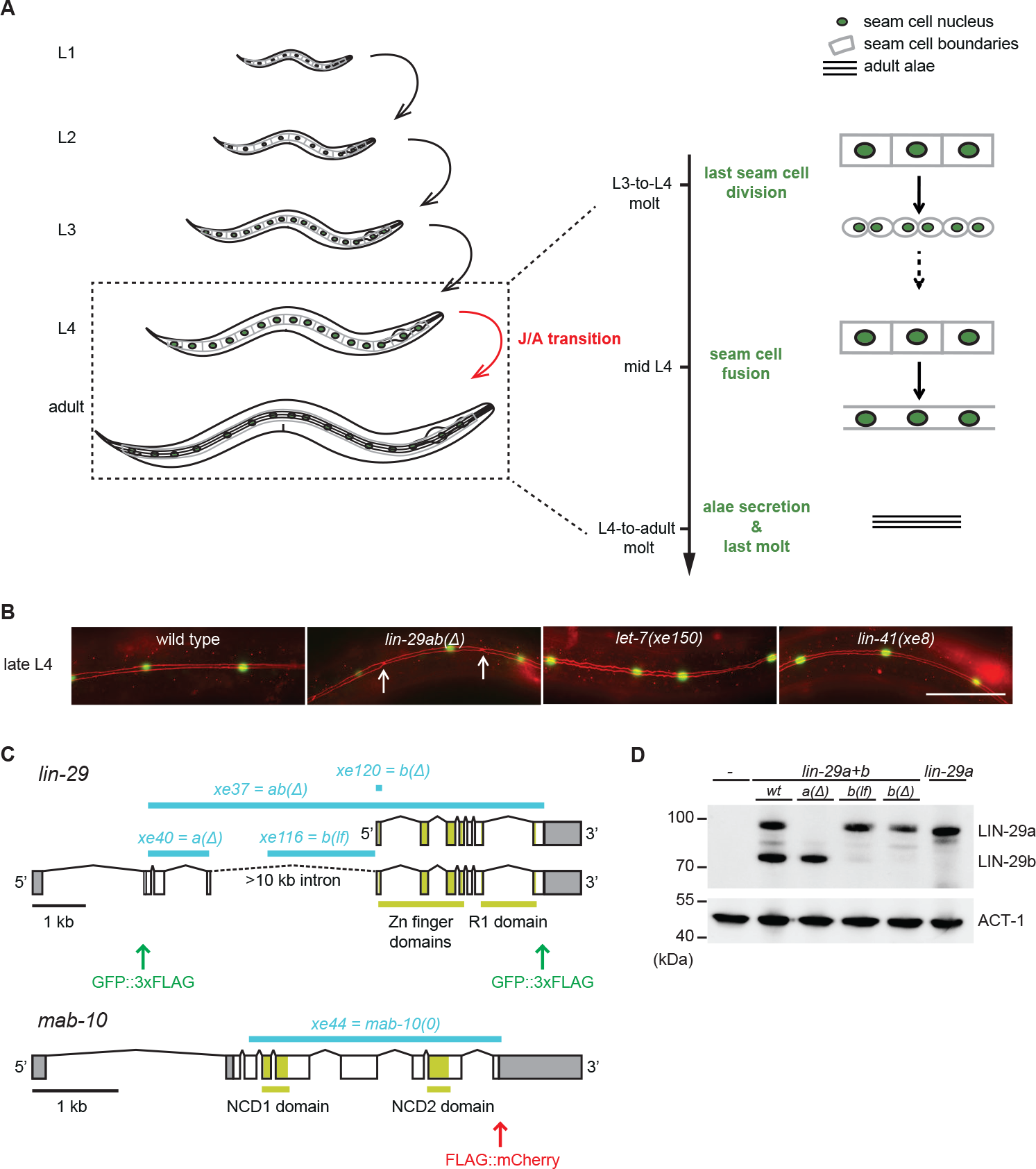
Uncoupling of coordinated execution of J/A transition events in *let-7* and *lin-41* mutant animals. A. Schematic representation of juvenile-to-adult (J/A) transition events in the *C. elegans* epidermis: final division of seam cells (square-shaped cells with green nuclei) at the L3-to-L4 molt; seam cell fusion into a syncytium during mid-L4 stage; synthesis of an adult cuticle containing lateral alae (three horizontal bars) at the L4-to-adult molt; and a subsequent exit from the molting cycle.
B. Micrographs of late L4-stage animals of indicated genotypes expressing *scm∷gfp* (*green*, marking seam cells) and *ajm-1∷mCherry* (*red*, marking hypodermal cell junctions). Arrows indicate cell boundaries between unfused cells. Representative of n>20. Scale bars: 50 μm.
C. Schematic representation of the *lin-29* and *mab-10* genomic regions. Mutant alleles and endogenously tagged alleles used in this study are indicated. A *gfp∷3×flag* towards the 5’ end specifically tags LIN-29a, while a *gfp∷3×flag* towards the 3’ end both isoforms at the shared C-terminus. Insertion of this C-terminal tag in a *lin-29(xe40[lin-29a(Δ)])* genetic background yields specific tagging of *lin-29b*. The Methods section provides a detailed explanation of *lin-29(xe120[lin-29b(Δ)])*. Allele numbers refer to modifications in otherwise wild-type backgrounds; numbers for equivalent mutations in the endogenously tagged backgrounds, used for protein detection by microscopy and Western blotting, are provided in Table S1.
D. Western blot of C-terminally GFP∷3×FLAG-tagged endogenous LIN-29a and LIN-29b proteins in the different mutant backgrounds using anti-FLAG antibody. Actin-1 is used as loading control.

Genetic screens have identified precocious and retarded mutations (Ambros and Horvitz, 1984), which cause animals to exhibit somatically adult features before reaching sexual maturity or retain juvenile somatic features after reaching sexual maturity, respectively. Among the factors thus identified, LIN-29, a transcription factor of the EGR/Krüppel family, is considered the downstream-most gene of the heterochronic pathway (Rougvie and Moss, 2013). Indeed, LIN-29 accumulates immediately prior to transition to adulthood during the last (L4) larval stage (Bettinger et al., 1996).

Current models of the heterochronic pathway depict a simple linear chain of events during the last larval stages that leads to upregulation of LIN-29 (Faunes and Larraín, 2016; Moss and Romer-Seibert, 2014; Rougvie and Moss, 2013). The miRNA *let-7* accumulates during the L3 stage to inhibit synthesis of the RNA binding protein LIN-41 (Ding and Großhans, 2009; Reinhart et al., 2000). Since LIN-41 translationally represses *lin-29* (Aeschimann et al., 2017), its decay during L4 allows for expression of LIN-29. As all four epidermal J/A events require LIN-29 (Ambros and Horvitz, 1984; Bettinger et al., 1997), this pathway architecture can ensure their coordinated execution.

However, multiple lines of evidence challenge this simple linear model. First, LIN-29 occurs in two protein isoforms, LIN-29a and LIN-29b (Rougvie and Ambros, 1995), and LIN-41 appears to silence only *lin-29a* but not *lin-29b* (Aeschimann et al., 2017). Second, *lin-41(0)* mutant precocious phenotypes, unlike the retarded *lin-29(0)* mutant phenotypes, are only partially penetrant (Slack et al., 2000), indicating additional control of LIN-29 beyond repression by LIN-41. Third, *let-7* mutations do not recapitulate all phenotypes of *lin-29(0)* (Ambros, 1989), as *let-7* appears dispensable for proper timing of seam cell fusion (Hunter et al., 2013).

Here we address these discrepancies by studying the regulation of the two LIN-29 isoforms and their functions in J/A transition. LIN-29a and LIN-29b share most of their protein sequence with the exception of 142 amino acids at the N-terminus that are unique to LIN-29a (Figure 1B). Previous studies have suggested that these isoforms function redundantly: they share a common co-factor, MAB-10 (Harris and Horvitz, 2011), they have similar expression patterns, and they are interchangeable in complementation analysis (Bettinger et al., 1996, 1997). However, employing isoform-specific mutations and endogenous protein tagging, we show here that the *lin-29a* and *lin-29b* isoforms differ in function and expression patterns. The most striking functional difference occurs in seam cell fusion, which relies only on LIN-29b, but not on LIN-29a or MAB-10. Whereas *lin-29a* is regulated by LIN-41, the *lin-29b* isoform is regulated by the transcription factor HBL-1, which results in a distinct spatiotemporal expression of the two isoforms. The expression patterns alone can explain the unique phenotypic consequences of the isoform-specific mutations, whereas the sequence difference in the N-terminal portion of the two isoforms does not contribute to functional differences. Coordination of the activities of LIN-29a and LIN-29b is achieved through LIN-28, an RNA-binding protein that regulates both HBL-1 and *let-7*–LIN-41. Taken together, our findings help to reframe the regulatory logic that enables the heterochronic pathway to coordinate different events into an overall larval-to-adult transition program.

## RESULTS

### Regulation of LIN-29 by *let-7* and LIN-41 is dispensible for triggering seam cell fusion

A model of simple linear control in the heterochronic pathway predicts that *lin-29(0)* mutations cause the same phenotypes as upstream mutations that impair the activation of *lin-29*, such as *let-7(0)*. Indeed, the phenotypes of the two mutant strains overlap extensively (Ambros and Horvitz, 1984; Bettinger et al., 1997; Reinhart et al., 2000). However, they are not identical: Whereas *lin-29(0)* mutant animals fail to execute seam cell fusion (Bettinger et al., 1997), this process was reported to occur normally in *let-7(mn112)* mutant animals (Hunter et al., 2013). To validate this unexpected observation, we examined a newly created *let-7* null mutant strain, *let-7(xe150)*, which lacks *let-7* expression due to deletion of the promoter (a gift from J. Kracmarova). Additionally, we used the temperature-sensitive *let-7(n2853)* strain (Reinhart et al., 2000) at the restrictive temperature, 25°C. Employing an *ajm-1∷mCherry* marker to visualize the seam cells boundaries, we observed that seam cells fused normally in 100 % of the *let-7(xe150)* and *let-7(n2853)* mutant animals, confirming that *let-7* is indeed dispensable for seam cell fusion (Figure 1B, Table S2). By contrast, complete loss of *lin-29* in *lin-29(xe37)* null mutant animals (Aeschimann et al., 2019) caused failure of seam cell fusion in all animals (Figure 1B).

Contrasting with the normal seam cell fusion in *let-7* mutant animals, overexpression of *lin-41* had previously been reported to prevent seam cell fusion (Slack et al., 2000). However, when we examined *lin-41(xe8[ΔLCS])* mutant animals (Figure 1B), which lack the *let-7* complementary sites (LCSs) in the *lin-41* 3’UTR and thus exhibit sustained high LIN-41 levels during the L4 stage (Aeschimann et al., 2017; Ecsedi et al., 2015), seam cell fusion occurred normally. Hence, the activity of the *let-7*–LIN-41 module cannot account for all the J/A transition events regulated by *lin-29*.

### Seam cell fusion requires LIN-29b but not its co-factor MAB-10 nor LIN-29a

Recently, we showed that *let-7*–LIN-41 regulate *lin-29a* (and *mab-10*), but possibly not *lin-29b* (Aeschimann et al., 2017, 2019). Accordingly, in *let-7(0)* and *lin-41(ΔLCS)* animals, LIN-29a and MAB-10 levels are expected to remain low in the L4 stage while LIN-29b can accumulate. Hence, we wondered whether LIN-29b would suffice for seam cell fusion and thus explain the phenotypic discrepancy between *let-7(0)* and *lin-29(0)* mutant animals. To test this possibility, we generated two *lin-29b* isoform-specific mutations (Methods). One, where we deleted the putative promoter of *lin-29b*, achieved extensive, but not complete depletion of LIN-29b while leaving LIN-29a levels unaltered (Figure 1C,D). We will refer to it as *lin-29b(lf).* The other, *lin-29b(Δ)*, where we altered *lin-29b* translation initiation to translate it out of frame, caused a complete loss of LIN-29b, but we cannot exclude a modest depletion of LIN-29a (Figure 1C,D). Thus, failure to see a phenotype of interest in the *lin-29b(Δ)* mutant excludes an essential function of the b isoform. Conversely, observation of a phenotype in the *lin-29b(lf)* mutant reveals an essential contribution of the b isoform to this phenotype, although we might under-estimate the extent of this contribution. We engineered each mutation into two different backgrounds, wild-type animals for functional studies, and animals containing a GFP∷3×FLAG-tag at the shared C-terminus of the LIN-29 isoforms for expression analysis by Western blotting and imaging (Table S1).

We used these, and the previously generated *lin-29(xe40)* (henceforth *lin-29a(Δ)*) and *mab-10(xe44)* (henceforth *mab-10(0)*) mutant strains (Aeschimann et al., 2019) to study the individual contributions of LIN-29a, LIN-29b and their co-factor MAB-10 to the different J/A transition events. MAB-10 itself is thought not to directly bind to DNA, but to bind to and modulate the activity of both LIN-29 isoforms (Harris and Horvitz, 2011). Consequently, in the LIN-29a- or LIN-29b-specific single mutants, either of the two LIN-29 isoforms left can still act together with its co-factor MAB-10, while double mutant animals lacking both MAB-10 and one LIN-29 isoform are left with only the other LIN-29 isoform, acting without the co-factor MAB-10.

To determine the individual roles of the LIN-29 isoforms in seam cell fusion, we used the *ajm-1∷mCherry* marker to count the number of unfused junctions in the late L4 stage, after the last seam cell division and before the last molt (Figure 2A-C). *lin-29a(Δ)* and *mab-10(0)* single mutant animals, as well as *mab-10(0) lin-29a(Δ)* double mutant animals showed unperturbed seam cell fusion. This is consistent with the functional fusion seen in *let-7(0)* and *lin-41(ΔLCS)* animals (Figure 1B).

**Figure 2:**
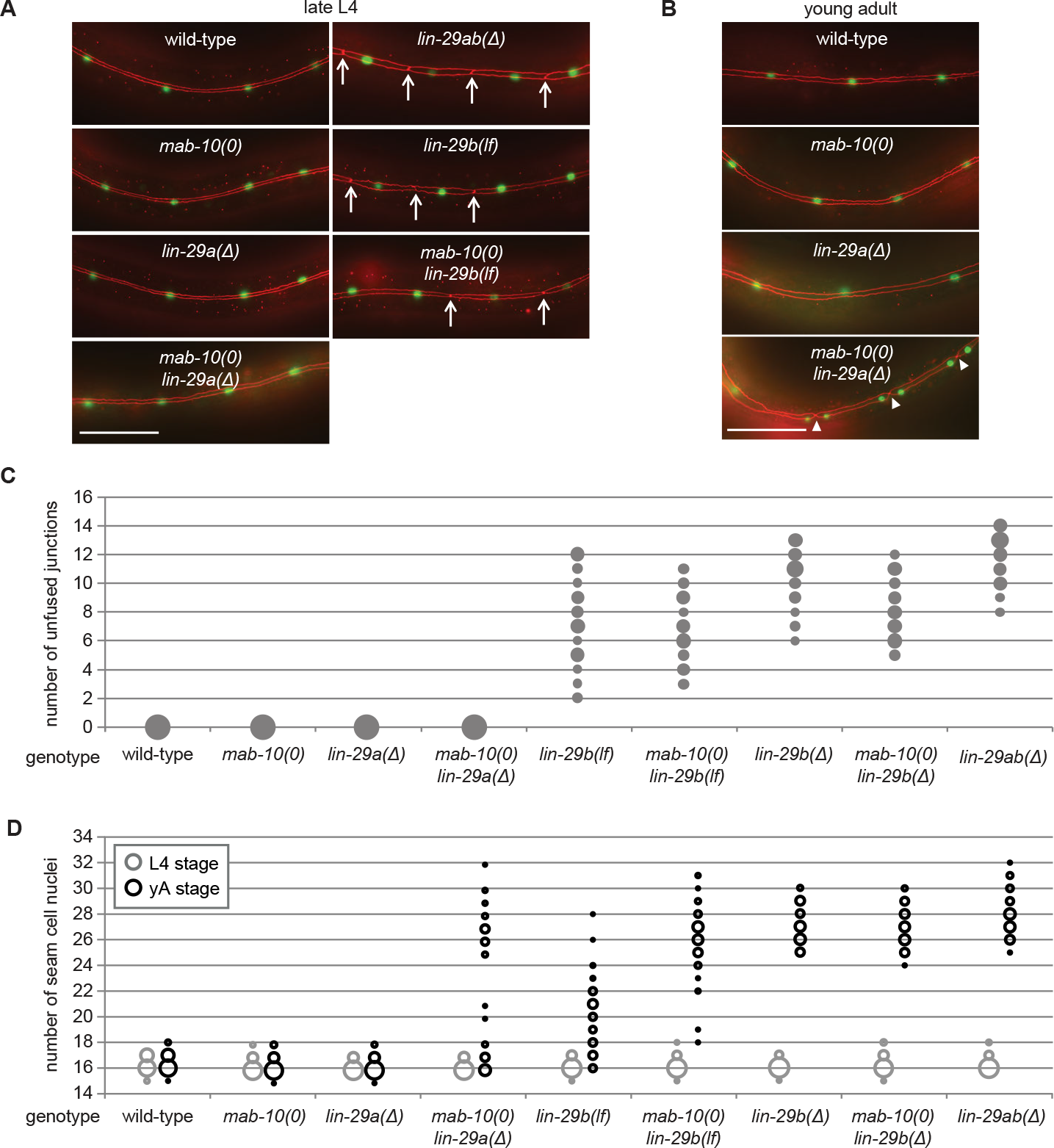
LIN-29b has a fundamental role in the regulation of early J/A transition events. (A-B) Micrographs of late L4 stage (A) and young adult (B) animals of indicated genotype expressing *scm∷gfp* (*green*, marking seam cells) and *ajm-1∷mCherry* (*red*, marking seam cell boundaries). Arrows indicate cell boundaries between unfused cells. Scale bars: 50 μm. Bubble chart showing quantification of unfused seam cell junctions of L4 larval stage animals of the indicated genetic backgrounds. Areas of bubbles represent the percentage of worms with the respective number of unfused junctions (n>20 for each genotype). Bubble chart showing seam cell numbers in L4 larval stage and young adult (yA) animals of the indicated genetic backgrounds. Areas of bubbles represent the percentage of worms with the respective number of seam cells (n=20 for L4, n>50 for yA worms per genotype).

In striking contrast to these findings, we observed penetrant seam cell fusion defects in both *lin-29b(lf)* and *lin-29b(Δ)* single mutant strains, and this phenotype was not enhanced by concomitant loss of *mab-10* expression, i.e., in *mab-10 lin-29b* double mutant animals. These findings support the notion that the *let-7*–LIN-41–LIN-29a/MAB-10 module is dispensable for seam cell fusion. They also reveal an unanticipated specialization of LIN-29 isoforms, with LIN-29b being both necessary and sufficient for wild-type seam cell fusion. We conclude that the two LIN-29 isoforms fulfill distinct and non-redundant functions.

### Fully functional cell cycle exit of seam cells requires both LIN-29 isoforms and their co-factor MAB-10

Prompted by the discovery of a non-redundant function of LIN-29 isoforms in seam cell fusion, we surveyed their individual contributions to other J/A transition events. First, we examined exit of seam cells from the cell cycle. As shown previously (Aeschimann et al., 2019; Ambros and Horvitz, 1984), *lin-29ab(Δ)* (*lin-29(xe37)*) animals exhibit a fully penetrant phenotype: seam cells do not exit the cell cycle but instead continue to divide so that all animals show at least 25 seam cells instead of the canonical 16 at the young adult stage (Figure 2D). By contrast, seam cells exit the cell cycle normally in *lin-29a(Δ)* and *mab-10(0)* animals. A partially penetrant defect occurs in *mab-10(0) lin-29a(Δ)* double mutant animals (Figure 2D), recapitulating the phenotype of *let-7(n2853)* and *lin-41(ΔLCS)* animals (Aeschimann et al., 2019). Unlike loss of LIN-29a alone, depletion of LIN-29b alone suffices to permit unscheduled seam cell divisions in young adult animals (Figure 2D). The phenotype is more penetrant in *lin-29b(Δ)* than in *lin-29b(lf)* animals. Moreover, loss of *mab-10* enhances the *lin-29b(lf)* but not the *lin-29b(Δ)* mutant phenotype.

Collectively, these data thus confirm an involvement of all three factors, LIN-29a, LIN-29b and MAB-10 in seam cell cell cycle exit. The data also suggest a more prominent role for LIN-29b versus LIN-29a, and a function for MAB-10 in promoting LIN-29 activity in this process.

### Uncoupling of nuclear division and differentiation programs in seam cells

Cell division and terminal differentiation are normally mutually exclusive, tightly coupled events. In fact, exit from the cell division cycle is frequently considered a central aspect of terminal cell differentiation (Myster and Duronio, 2000). Hence, we were surprised to find that nuclear divisions continued to occur after seam cells had fused at the L4 stage in *mab-10(0) lin-29a(Δ)* mutant animals (Figure 2A-D). We investigated this apparent discrepancy further and found that after normal fusion in the L4 stage, cell junctions re-form after additional nuclear divisions in the adult stage. This resulted in the appearance of pairs of seam cell nuclei, separated from other pairs by cell junctions. In this case cell boundaries surround pairs of ‘cousins’ rather than individual sister cell nuclei (Figure 2B). We conclude that exit from the cell cycle is not elicited by cell fusion, and vice versa, that cell fusion does not require permanent cell cycle exit.

### Both LIN-29 isoforms have important but distinct functions in alae formation

To understand LIN-29 isoform function in late J/A transition events, we examined the cuticles of young adults of the different genetic mutant backgrounds by DIC microscopy. An adult cuticle is characterized by alae (Figure 3A(I)), ridges along the lateral sides of the whole worm that are secreted by seam cells (Singh and Sulston, 1978). Consistent with previous results (Ambros and Horvitz, 1984), we observed a complete lack of alae in *lin-29ab(Δ)* young adults (Figure 3B). *lin-29a(Δ)* mutant animals were partially defective in alae formation, exhibiting weak alae structures (Figure 3A(II)) that either covered the whole worm (“complete”) or at least 50 % of the body length (“partial”) (Figure 3B). Animals depleted of LIN-29b either exhibited only alae patches or lacked alae entirely (Figure 3A(III, IV)). *lin-29b(Δ)* animals had a more penetrant “no alae” phenotype than *lin-29b(lf)* mutant animals, with almost half of the former animals lacking any detectable alae structure (Figure 3B). However, neither *lin-29b* mutation fully recapitulated the *lin-29ab(Δ)* phenotype, and the alae defects were qualitatively distinct in the *lin-29a(Δ)* and the *lin-29b(Δ)* mutant animals. Finally, *mab-10* mutant animals displayed normal, wild-type alae formation (Harris and Horvitz, 2011), and absence of MAB-10 did not enhance the phenotypes of any of the *lin-29a* or *lin-29b* mutations (Figure 3B).

**Figure 3:**
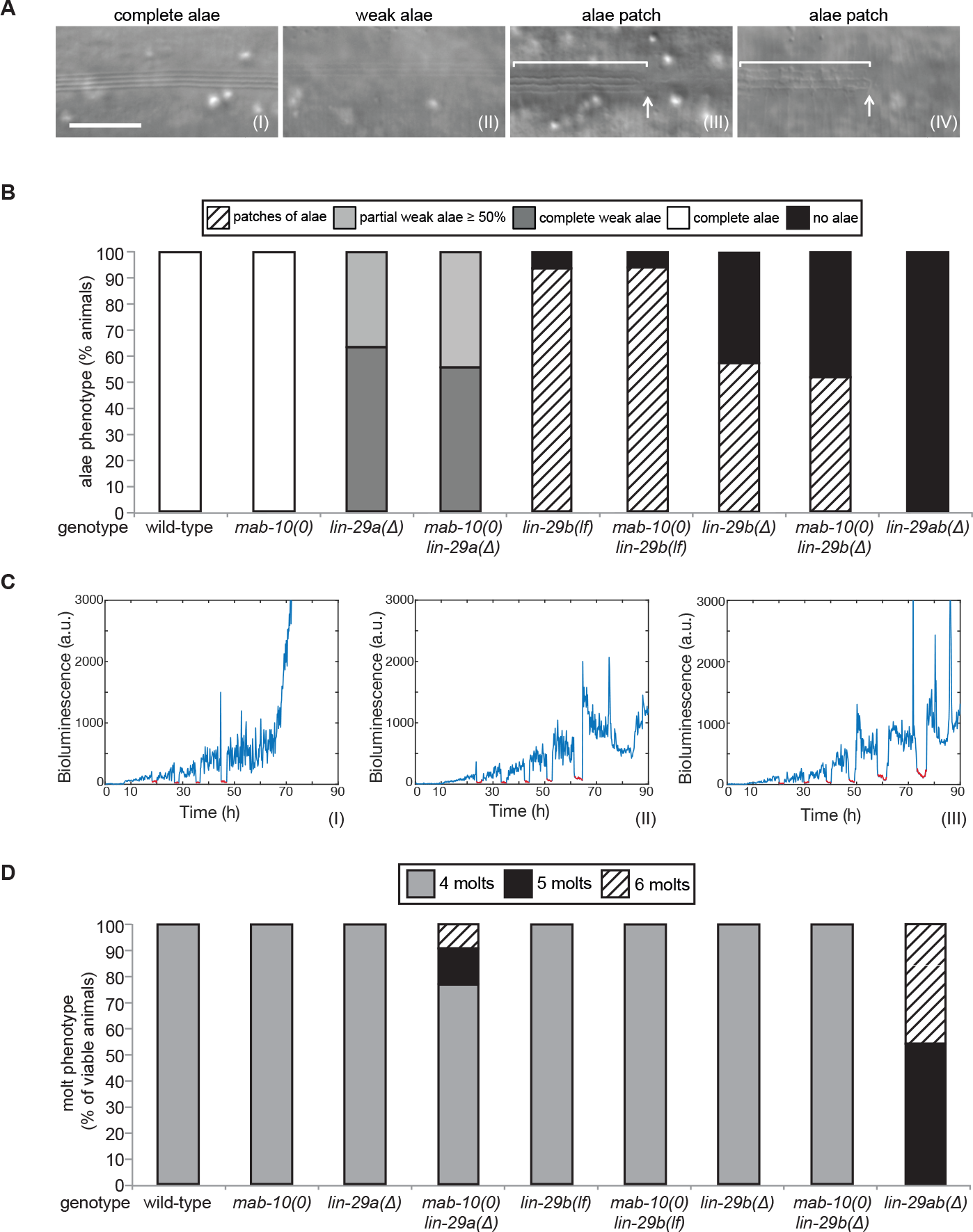
Late J/A transition events are regulated by both LIN-29 isoforms. A. Micrographs illustrating categories of alae structures observed on the cuticle of wild-type (I) and mutant (II – IV) young adult animals. The example pictures show the weak alae phenotype of a (II) *mab-10(0) lin-29a(Δ)* mutant animals, and alae patches of (III) *mab-10(0) lin-29b(lf)* and (IV) *mab-10(0) lin-29b(Δ)* animals, respectively. Scale bars: 10 μm (100X magnification).
B. Quantification of different alae structures of young adult worms of indicated genotypes (n>30).
C. Examples of luciferase assay traces revealing four (I), five (II) or six (III) molts through a drop in luciferase signal (red segment). Examples are from wild-type (I) and *lin-29ab(Δ)* (II-III).
D. Quantification of abnormal number of molts of indicated genotypes (n>20) based on assay shown in C. A small fraction of *mab-10(0) lin-29a(Δ)* animals die at the J/A transition (Aeschimann et al., 2019); these animals were censored and not included in the quantification.

We conclude that both LIN-29a and LIN-29b, but not MAB-10, are required for wild-type alae formation, and that their functions are partially distinct.

### LIN-29a and LIN-29b function redundantly to regulate the exit from the molting cycle

The last event of the J/A transition that we examined was molting. Using a high-throughput assay (Meeuse et al., 2019; Olmedo et al., 2015), we counted molts in >20 animals for each genotype. We confirmed a previous report (Ambros and Horvitz, 1984) that *lin-29* was required for the exit from the molting cycle by observing that all *lin-29ab(Δ)* animals exhibited at least one extra molt, with ~50% exhibiting two extra molts. By contrast, animals with almost all other single or double mutant combinations, i.e., *mab-10(0), lin-29a(Δ), lin-29b(lf), lin-29b(Δ)* single mutations and *mab-10(0) lin-29b(lf)* and *mab-10(0) lin-29b(Δ)* double mutations, did not display defects in exiting the molting cycle. The only exception were *mab-10(0*) *lin-29a(Δ)* animals, of which a small percentage executed one or two extra molts (Figure 3D).

We conclude that LIN-29a and LIN-29b have a redundant function in promoting the exit from the molting cycle. Their co-factor MAB-10 has a minor contribution that becomes detectable in a sensitized background.

### Functional differences do not result from difference in molecular sequence

Since our experiments show that LIN-29a and LIN-29b have partially distinct functions (Figure S1), we asked whether their molecular differences, i.e., the unique N-terminal extension of 142 amino acids of LIN-29a (Figure 1C), are responsible for functional specialization. To address this issue, we genetically engineered the *lin-29* locus such that this N-terminal extension was removed by deleting the *lin-29a*-specific coding exons (exons 2-4). This mutation created a *lin-29a* transcript identical to that of *lin-29b* except for harboring the *lin-29a* 5’UTR and encoding 23 additional N-terminal amino acids. We confirmed by Western blotting that this truncated LIN-29a(ΔN) protein had a similar size to LIN-29b and accumulated at roughly wild-type LIN-29a levels (Figure 4A). Moreover, the removal of the N-terminus did not affect the spatiotemporal expression pattern of *lin-29a*, and its expression remained under control of LIN-41 (Figure S2C-S2D).

**Figure 4:**
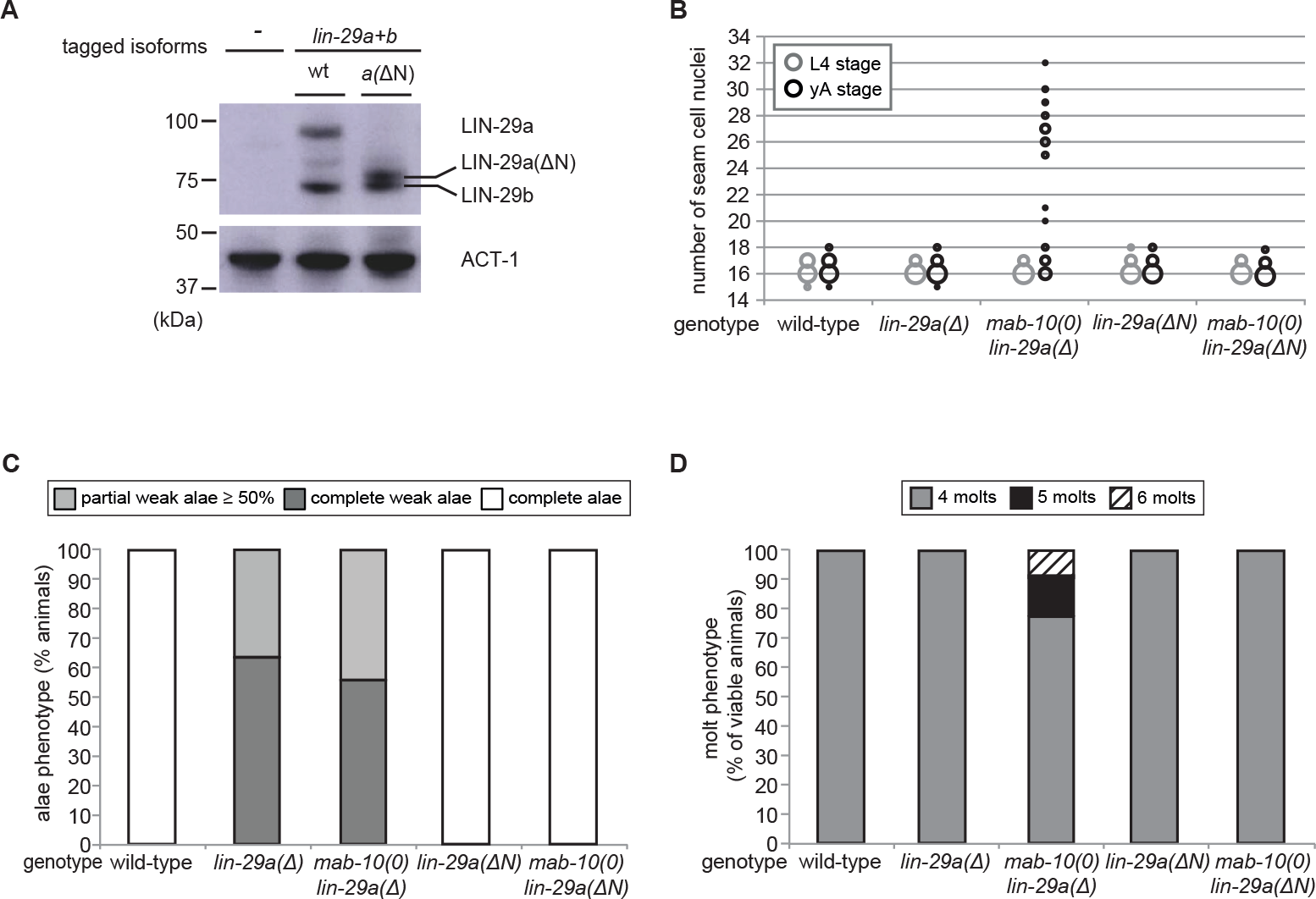
The LIN-29a-specific domain is dispensable for the execution of the J/A transition. A. Western blot of endogenous C-terminally GFP∷3×FLAG-tagged LIN-29a and LIN-29b proteins in the *lin-29a(ΔN)* background (HW2408) using anti-FLAG antibody. Actin-1 is used as loading control.
B. Seam cell number quantification in L4 larval stage and young adult (yA) animals of the indicated genetic backgrounds. Areas of bubbles represent the percentage of worms with the respective number of seam cells (n>25 for L4, n>25 for yA worms per genotype). The data for *lin-29a(Δ)* and *mab-10(0) lin-29a(Δ)* is re-plotted from Figure 2 for the purpose of comparison.
C. Quantification of different alae structures of young adult worms of indicated genotypes (n>20). The data for *lin-29a(Δ)* and *mab-10(0) lin-29a(Δ)* is re-plotted from Figure 3 for the purpose of comparison.
D. Quantification of abnormal number of molts of indicated genotypes (n>20). The data for *lin-29a(Δ)* and *mab-10(0) lin-29a(Δ)* is re-plotted from Figure 3 for the purpose of comparison.

We characterized the phenotypes of *lin-29a(ΔN)*, alone or in combination with a *mab-10(0)* allele, for all four J/A transition events. In all assays, deletion of the LIN-29a-specific N-terminus did not compromise LIN-29a function, resulting in a wild-type phenotype for all four J/A transition events (Figure 4B-D, Figure S2). This data suggests that the distinct functions of LIN-29a and LIN-29b are not mediated by sequence differences, although we cannot formally exclude a contribution of the remaining unique 23 amino acids.

### *lin-29a* and *lin-29b* differ in spatiotemporal expression patterns in the epidermis

Given the apparent absence of differences in molecular function between the two LIN-29 isoforms, we revisited their developmental expression patterns. *lin-29a* and *lin-29b* transcripts were previously reported to exhibit largely similar temporal patterns of accumulation, being both detectable from L1 stage on, with increasing levels during development, peaking at the L4 stage, and decreasing in adulthood (Rougvie and Ambros, 1995). Furthermore, a similar spatiotemporal expression pattern was deduced from promoter activity reporter experiments (Bettinger et al., 1996). However, transcript quantification and promoter activity measurements do not account for additional layers of post-transcriptional regulation, such as LIN-41-mediated translational repression of *lin-29a* (Aeschimann 2017). Previous analysis of LIN-29 protein accumulation by immunofluorescence could not distinguish between the two protein isoforms (Bettinger et al., 1996).

To elucidate the temporal expression pattern of *lin-29* isoforms on the protein level, we examined animals carrying a C-terminal GFP∷3×FLAG tag that marks both isoforms. Using Western blotting, we could distinguish the two isoforms by their distinct sizes. The levels of the LIN-29b protein agreed with its previously reported pattern of mRNA accumulation, with detectable accumulation throughout larval stages and an increase in levels in the L3 and L4 stages (Figure 5A). By contrast, although the *lin-29a* transcript is detectable in L1 and abundant in L2 (Rougvie and Ambros, 1995), LIN-29a protein was undetectable in L1 or L2 stage worms, accumulated weakly in L3 stage, and peaked in L4 stage worms. This is consistent with its post-transcriptional regulation by LIN-41 (Aeschimann et al., 2017). We note that in late larval stages and adults, we also detect a third band migrating in between the LIN-29a and LIN-29b bands (Figure 1D, 5A asterisk). We did not find evidence for a transcript encoding this intermediate-size protein in available paired-end gene expression data (F. Gypas, personal communication) and the molecular nature of this band remains unclear. For the purpose of this study, we considered only the two previously described isoforms (Rougvie and Ambros, 1995).

**Figure 5:**
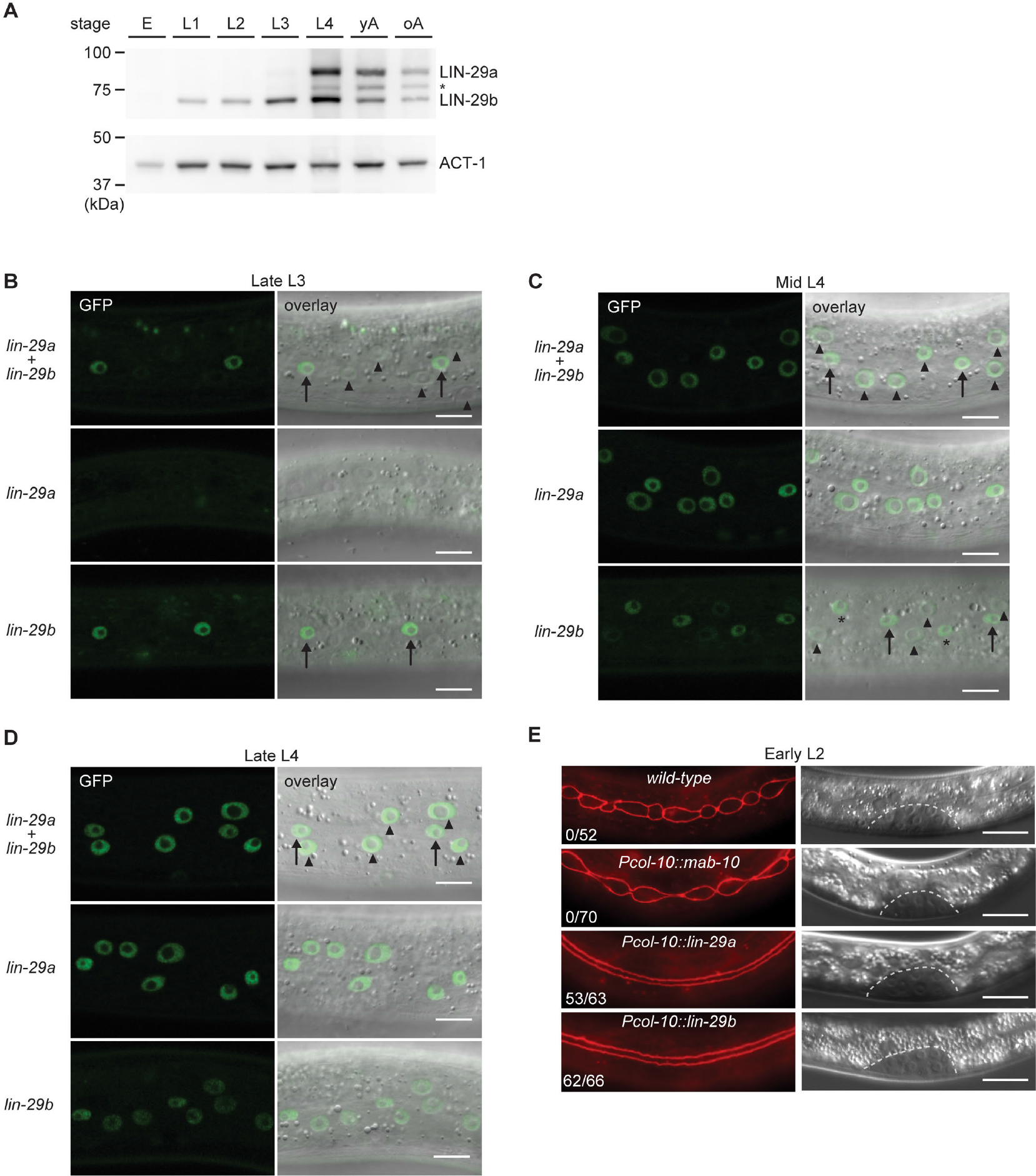
Spatial and temporal expression pattern of LIN-29a and LIN-29b. (A) Western blot of C-terminally GFP∷3×FLAG-tagged endogenous LIN-29a and LIN-29b proteins in different developmental stages using anti-FLAG antibody. Actin-1 is used as loading control. The asterisk indicates a putative degradation product. (B-D) Confocal images of endogenously tagged LIN-29 protein isoforms in the epidermis of animals at the indicated developmental stages. Animals were staged by examination of gonad development as late L3 stage (B), and mid- (C) and late (D) L4 stage. Arrows indicate seam cell, arrowheads hyp7 nuclei. Scale bars: 10 μm. (E) Micrographs of early L2 animals expressing the indicated transgenes (‘wild-type’: no transgene control) and *ajm-1∷mCherry* (*red*, marking hypodermal cell junctions). Animal stage was confirmed by gonad morphology as shown in the DIC pictures. Numbers indicate fractions of animals with complete precocious seam cells fusion. Scale bars: 10 μm.

To examine the spatiotemporal expression patterns of the two *lin-29* isoforms *in vivo*, we utilized strains with GFP∷3×FLAG tags on the individual, endogenous LIN-29 isoforms. For *lin-29a*, we used a published strain carrying a GFP∷3×FLAG tag at the unique N-terminus of LIN-29a (Aeschimann et al., 2019). These animals did not display any overt phenotypes. Since we could not create a LIN-29b-specific tag due to lack of isoform-specific sequence, we used a C-terminal fusion that tags both isoforms and mutated the *lin-29a(Δ)* isoform in this background (Figures 1D). Hence, we study *lin-29b* expression in a *lin-29a(Δ)* mutant background. The resulting strain exhibited an increased occurrence of protruding vulva (Pvl) and vulval bursting phenotypes relative to *lin-29a(Δ)* animals without the tag (data not shown) (Aeschimann et al., 2019), indicating that the C-terminal tag may partially impair LIN-29 protein function.

Using live imaging of the GFP-tagged proteins, we observed LIN-29 isoform accumulation. In agreement with immunofluorescence-based analysis (Bettinger et al., 1996), we detected signal in the epidermis, but also in other regions such as the head, the tail and the vulva (Figure 5B-D, Figure S3). Considering our interest in the role of LIN-29 in the epidermal J/A transition, we focused on the detailed characterization of LIN-29a and LIN-29b accumulation in the skin. LIN-29b was first detectable in lateral seam cell nuclei of late L3 worms (Figure 5B). Subsequently, in mid-L4 stage animals, it also accumulated, weakly, in the major hypodermal syncytium hyp7 (Figure 5C). By contrast, LIN-29a was not detected in either seam cells or the major hypodermal syncytium hyp7 during the L3 stage (Figure 5B). Instead, LIN-29a accumulation began in both cell types in mid-L4 stage worms. (Figure 5C). Finally, protein levels of both LIN-29a and LIN-29b peaked in lateral seam cells and hyp7 in late L4 stage animals and persisted into adulthood (Figure 5D, data not shown).

Taken together, our detailed analysis shows that *lin-29a* and *lin-29b* differ in their spatial as well as temporal expression patterns in the epidermis, likely explaining their distinct function in the four J/A transition events.

### Precocious expression of either *lin-29a* or *lin-29b* can induce seam cell fusion

To formally test if the functional difference between the two *lin-29* isoforms stems from their distinct expression patterns rather than molecular differences, we sought to alter the patterns experimentally. To this end, we expressed either *lin-29a* or *lin-29b* from a single-copy transgene under the control of the epidermal *col-10* promoter, which is active from the first larval stage onwards. To avoid potential toxic effects of temporal misexpression of *lin-29* in early larval stages, we added an auxin-inducible degron (AID) tag (Zhang et al., 2015) to the transgenic LIN-29 isoforms and prevented LIN-29 protein accumulation by addition of auxin to maintain the transgenic lines (Methods). We then let animals hatch in the absence of auxin and plated starved, synchronized L1 stage larvae on food to permit LIN-29 protein accumulation, before we examined seam cell fusion, the J/A transition event that relies exclusively on LIN-29b. Strikingly, *col-10*-driven expression of either isoform induced precocious seam cell fusion. Specifically, we observed individual animals with fused seam cells already at 10 hours after plating at 25°C, i.e., during the L1 stage. At 15 hours after plating, in early L2, all animals had at least partially fused seam cells, with >80% of animals with complete fusion (Figure 5E). By contrast, expression of *mab-10* from the *col-10* promoter did not cause any precocious seam cell fusion at this time (Figure 5E). We conclude that LIN-29 upregulation alone is sufficient to trigger fusion of seam cells, and that upregulation of either LIN-29 isoforms suffices. This data further validates that the LIN-29b-specific function observed in seam cell fusion under physiological conditions (Figure 2A-2C) is not due to sequence differences between the two isoforms, but due to differences in their expression.

### Distinct regulation of *lin-29* isoforms through LIN-41 and HBL-1

We wondered how the distinct expression patterns of *lin-29a* and *lin-29b* are generated. We have previously shown that *lin-29a* is translationally regulated by LIN-41 (Aeschimann 2017). However, the regulation of *lin-29b* expression is poorly understood. We noticed that the alae phenotype seen in *lin-29b* mutant adults, small patches of well-formed alae, correspond to that seen one stage earlier in precocious *lin-41(0)* mutant animals (Abrahante et al., 2003). By contrast, the weaker but longer patches of alae seen in *lin-29a(Δ)* mutant adults are reminiscent of the appearance of precociously formed alae in HBL-1-depleted L4 stage larvae (Abrahante et al., 2003). Hence, we wondered whether HBL-1, a transcription factor of the Hunchback/Ikaros family (Abrahante et al., 2003; Lin et al., 2003), and LIN-41 might act on distinct *lin-29* isoforms.

Indeed, when we examined expression of *lin-29* at the early L3 stage using the isoform-specific GFP-tags on LIN-29, we found that, consistent with earlier findings (Aeschimann et al., 2017), LIN-41 depletion caused precocious accumulation of LIN-29a but not LIN-29b (Figure 6A-B). By contrast, early L3 stage animals exposed to *hbl-1* RNAi displayed precocious accumulation of LIN-29b but not LIN-29a (Figure 6A, C). Of note, in contrast to LIN-29a accumulation upon LIN-41 depletion, this LIN-29b accumulation occurred preferentially in the lateral seam cells, not hyp7 (Figure 6A). Finally, consistent with earlier evidence (Aeschimann et al., 2017), we observed precocious accumulation of a partially functional MAB-10∷mCherry protein (Pereira et al., 2019) in L3 stage animals depleted of LIN-41 in both seam cells and hyp7, whereas depletion of HBL-1 had no effect (Figure 6D).

**Figure 6:**
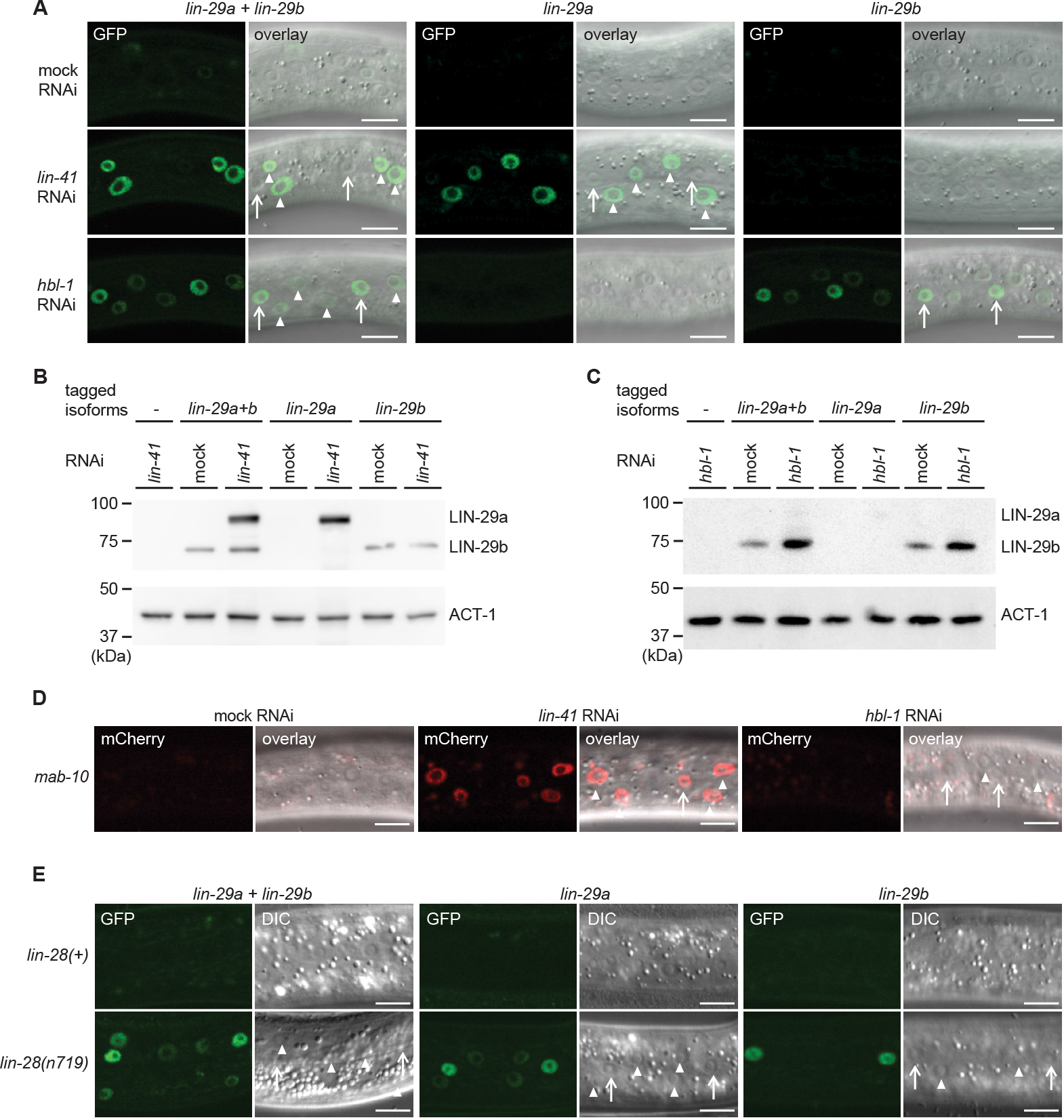
*lin-29a* and *lin-29b* expression are specifically regulated by LIN41 and HBL-1, respectively. (A) Confocal images of endogenously tagged LIN-29 protein isoforms in the epidermis (strains HW1822, HW1826, HW1835). Animals were grown at 25°C for 20 h on *lin-41, hbl-1* or mock RNAi bacteria. Arrows indicate seam cell, arrowheads hyp7 nuclei. Scale bars: 10 μm. (B-C) Western blot of GFP∷3×FLAG-tagged endogenous LIN-29a and LIN-29b proteins using anti-FLAG antibody. Actin-1 is used as loading control. Animals were grown for 20 h at 25°C to the L3 stage on mock RNAi and *lin-41* RNAi bacteria, respectively (B) or mock RNAi and *hbl-1* RNAi bacteria, respectively (C). Confocal images of endogenously tagged MAB-10 protein in the epidermis. Animals were grown at 25°C for 20 h on *lin-41, hbl-1* or mock RNAi bacteria. Arrowheads indicate hyp7 cells and arrows indicate seam cells. Scale bars: 10 μm. Confocal images of endogenously tagged LIN-29 protein isoforms in wild type or *lin-28(n719)* background in the epidermis (strains HW1822, HW1826, HW1835, HW1924, HW1925, HW1926). Animals were grown at 25°C for 20 h (control strains) and 22h (*lin-28(n719)* strains) to reach an equivalent developmental stage. Arrows indicate seam cell, arrowheads hyp7 nuclei. Scale bars: 10 μm.

We conclude that HBL-1 but not LIN-41 regulates LIN-29b accumulation. Conversely, MAB-10 and LIN-29a accumulation are regulated by LIN-41 but not by HBL-1. Whether the regulation of LIN-29b by HBL-1 is direct remains to be determined.

### LIN-28 regulates both *lin-29a* and *lin-29b*

The distinct function and regulation of *lin-29* isoforms that we observed reveals that the heterochronic pathway is not linear but branches into two parallel arms, LIN-41-LIN-29a/MAB-10 and HBL-1-LIN-29b. We wondered where in the pathway this bifurcation occurs. We have previously shown that *lin-41* is the only relevant target of the miRNA *let-7* (Ecsedi 2015, Aeschimann 2017, 2019), placing *let-7* into only one arm of the pathway. The heterochronic gene immediately upstream of *let-7* is *lin-28*, which directly inhibits *let-7* biogenesis (Lehrbach et al., 2009; Van Wynsberghe et al., 2011). Interestingly, *lin-28* was also reported to directly and positively regulate *hbl-1* (Vadla et al., 2012), making it a good candidate for being immediately upstream of the pathway’s branching point. Hence, we analyzed precocious expression of the LIN-29 protein isoforms in a putative null mutant background of *lin-28*, *lin-28(n719)*, at the early L3 stage. We observed that LIN-29a accumulates precociously in both hyp7 and lateral seam cells, and LIN-29b in seam cells of *lin-28* mutant animals. Hence, these data support our hypothesis that heterochronic pathway bifurcation occurs downstream of *lin-28*.

## DISCUSSION

In the more than three decades since the discovery of the first heterochronic genes (Ambros and Horvitz, 1984; Chalfie et al., 1981), accumulating genetic and molecular interaction data have facilitated the formulation of ever more refined heterochronic pathway models. A unifying concept of current heterochronic pathway models is that *C. elegans* J/A transition is triggered through a linear chain of events: *let-7* silences LIN-41, and thereby relieves *lin-29* from repression by LIN-41 (Ambros, 2011; Faunes and Larraín, 2016; Rougvie and Moss, 2013). The findings that we have presented here reveal that this concept requires revision. We demonstrate that LIN-29 occurs in two functionally distinct isoforms (Figure 7A) and that the functional separation is the result of differences in spatiotemporal expression patterns (Figure 7B) rather than in protein sequence. Unique expression patterns of the isoforms are the result of isoform-specific regulation: whereas *lin-29a* is under direct translation control by LIN-41 (Aeschimann et al., 2017), *lin-29b* is repressed, directly or indirectly, by the transcription factor HBL-1 (Figure 7C). Hence, J/A transition relies on two separate arms of the heterochronic pathway rather than a linear chain of events.

**Figure 7:**
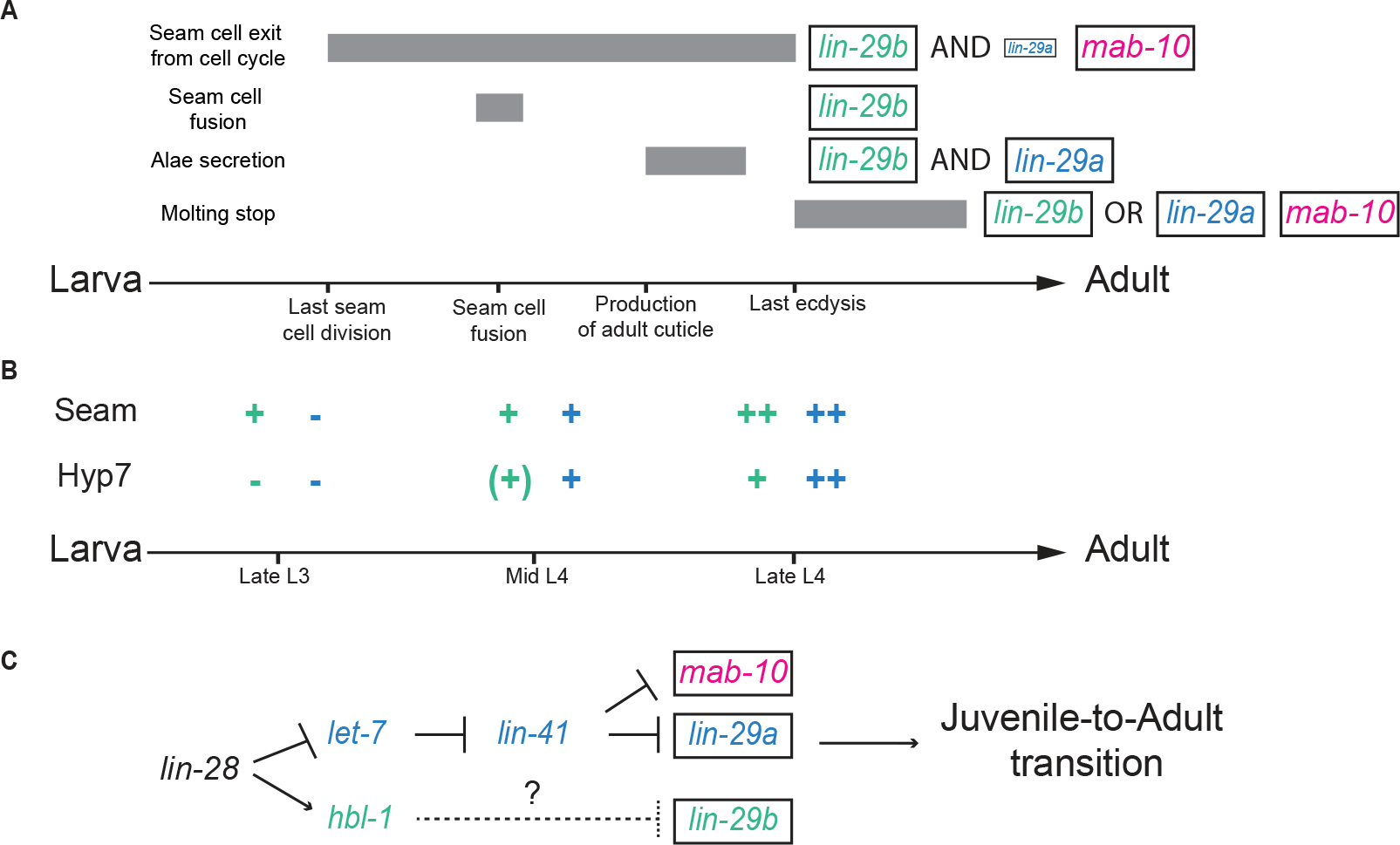
Summary. A. Illustration of the contribution of LIN-29 isoforms and the co-factor MAB-10 to the different J/A transition events. The arrow represents developmental time with relevant events indicated. Filled boxes indicate the duration of a specific J/A transition events. The exact time point when seam cells exit the cell cycle is unknown and might occur at any time after the last division at the L3/L4 molt and before the L4/adult molt. Although adults do not molt, it is unknown when exit from the molting cycle occurs.
B. Schematic depiction *of lin-29a* (*blue*) and *lin-29b* (*green*) expression patterns in seam cells and hyp7. Relevant developmental stages are indicated on the arrow representing developmental time.
C. Revised model of the heterochronic pathway. Two parallel arms of the heterochronic pathway exert their functions through distinct LIN-29 isoforms. Their activities are coordinated throught the upstream function of LIN-28.

The RNA-binding protein LIN-28 regulates both *lin-29* isoforms and thus coordinates their activity. Indeed, we propose that *lin-28* functions as the downstream-most heterochronic gene before branching, consistent with its dual functions of post-transcriptional activation of *hbl-1* expression (Vadla et al., 2012), and inhibition of *let-7* biogenesis (Lehrbach et al., 2009; Van Wynsberghe et al., 2011), which shields *lin-41* from repression by *let-7*.

Our findings and the revised heterochronic pathway model that we propose (Figure 7C) finally provide a mechanistic explanation for the previously reported redundancies between *lin-41* and *hbl-1* (Abrahante et al., 2003). However, they appear contradicted by other claims and findings from the published literature. First, the functional redundancy of LIN-29 isoforms inferred previously (Bettinger et al., 1997) disagrees with our finding of LIN-29 isoform specialization. Second, *lin-41* was described to affect seam cell fusion in both loss-of-function (Großhans et al., 2005) and over-expression experiments (Slack et al., 2000). This contrasts with a previous report that the *lin-41* regulator *let-7* was dispensable for seam cell fusion (Hunter et al., 2013) and, more explicitly, our conclusion that seam cell fusion relies on HBL-1–LIN-29b but not on *let-7*–LIN-41–LIN-29a. Finally, the new model, just like previous ones, fails to accommodate the apparent dual and antagonistic functions of HBL-1, namely suppression of adult cell fates in larvae, and suppression of larval cell fates in adults (Lin et al., 2003).

However, closer examination reveals that the published data are fully compatible with the new model. First, the conclusion that *lin-29b* is functionally equivalent to *lin-29a* was based on complementation of *lin-29(0)* mutant heterochronic phenotypes by a *lin-29b* transgene. In these experiments, *lin-29b* was expressed from a multicopy array (Bettinger et al., 1997) and thus likely over-expressed. Hence, rather than contradicting our model, these data provide further support for our conclusion that it is *lin-29* isoform expression, not protein sequence, that determines functional distinctions.

Second, although *lin29a* is dispensable for seam cell fusion in a wild-type context, our experiments (Figure 5) clearly demonstrate that, if expressed sufficiently early, *lin-29a* can induce seam cell fusion. This may precisely be the scenario in LIN-41 depleted animals, where LIN-29a (and MAB-10) accumulate precociously (Figure 6A, B, D and (Aeschimann et al., 2017)), thereby causing partial precocious seam cell fusion (Großhans et al., 2005).

Seam cell fusion failure in *lin-41* overexpression animals was inferred from the presence of seam cell junctions at the L4/adult molt, rather than by direct observation of syncytium formation in mid-L4 (Slack et al., 2000). However, the phenotype seems readily explained by re-appearance of junctions following a failure of syncytial nuclei to exit the division cycle. This is consistent with both normal fusion in a *lin-41(ΔLCS)* gain-of-function animals (Figure 1B) and *mab-10(0) lin-29a(Δ)* mutant animals, which extensively phenocopy *lin-41* gain-of-function mutant animals ((Aeschimann et al., 2019); Figure 2A-2C).

Third, we propose that a similar scenario of intrasyncytial cell divisions following seam cell fusion also explains the perplexing observation that adult *hbl-1* mutant adults exhibit seam cell junctions and a greater than wild-type number of seam cell nuclei (Lin et al., 2003), which, at the time, was interpreted as a retarded heterochronic phenotype. However, as we find that only LIN-29b accumulates precociously in HBL-1-depleted animals, we would expect them to undergo only a partial J/A transition at the L3 stage. Specifically, although seam cells fuse into a syncytium during L3, syncytial nuclei divide again in L4 and remain trapped in the seam (Abrahante et al., 2003). Hence, we propose that *hbl-1* mutant animals exhibit a purely precocious phenotype and that extra nuclei and seam cell junctions observed in *hbl-1* mutant adults are a consequence of an incomplete precocious J/A transition at the L3/L4 stage rather than reflection of a distinct, larval-fate suppressing function of HBL-1 in adults. Taken together, we find that the new model parsimoniously explains previously published and the present new data, and provides explanations for previously unaccounted phenotypes.

Our and the previous work (Abrahante et al., 2003; Lin et al., 2003) reveal a striking loss of coordination of J/A transition events in both HBL-1-depleted larvae and *mab-10(0) lin-29a(Δ)* double mutant adults. Specifically, in both cases, seam cells differentiate, as evidenced by fusion and alae formation, but fail to arrest the cell division cycle. This reveals that in seam cells, cell cycle arrest is not necessary for differentiation; i.e., proliferation and differentiation are not simply antagonistic processes. This finding highlights a need for factors that coordinate the activities of the two arms of the heterochronic pathway, and thus overall J/A transition. We have identified LIN-28 as a relevant factor, but others may exist. Nonetheless, coordination of J/A transition events through upstream activities of the heterochronic pathway rather than the previously proposed ‘terminal’ *let-7*–LIN-41–LIN-29 axis would seem to entail a surprising lack of robustness, making the pathway vulnerable to perturbations that can uncouple individual J/A transition events. Hence, we will be curious to learn whether this architecture is owed to other, unknown constraints or evolutionary history, or whether it has a particular benefit for the animal. At any rate, we speculate that additional layers of regulation will facilitate coordinated execution of the J/A transition.

Accumulation of LIN-29 has long been considered a key event for triggering J/A transition. We support this key function of LIN-29 in J/A transition by showing that LIN-29 accumulation alone is sufficient to trigger seam cell fusion as early as the L1 stage. While opposite temporal transformation phenotypes for loss-of-function and gain-of-function alleles are considered a hallmark of core heterochronic genes (Moss and Romer-Seibert, 2014), rather surprisingly, this had not previously been shown for LIN-29, and it had indeed been argued that LIN-29 accumulation was insufficient to trigger precocious J/A transition (Slack et al., 2000). However, our results are further supported by recent molecular evidence showing that *lin-29* is both necessary and sufficient for expression of adult collagens (Abete-Luzi and Eisenmann, 2018).

We propose that, in the unperturbed system, differences in LIN-29 dose sensitivity of the individual epidermal J/A transition events together with spatiotemporal differences in *lin-29* isoform expression drive their temporal order (Figure 7A-7B). Specifically, we hypothesize that seam cell fusion rely on LIN-29 accumulation in only the seam, explaining its early occurrence and dependence on only LIN-29b, which accumulates early in this tissue. By contrast, accumulation of LIN-29b alone appears insufficient to drive cessation of seam cell division cycles. Although the last seam cell division normally happens at the L3-to-L4 molt, it is unclear when these cells exit the cell cycle (Figure 7A). In fact, our data suggest the possibility of a gradual effect where LIN-29a and LIN-29b can, individually, and more extensively jointly, delay division, i.e., slow down the cell cycle, but where terminating it requires full LIN-29 activity, and thus MAB-10. Consistent with this notion, exit from the mitotic cycle is the only epidermal J/A transition event for which we observed a defect in *mab-10(0)* single mutant animals, but the additional divisions occurred much later than in the other mutant combinations (data not shown). We note that others reported also extra molts in, mostly older and male, *mab-10* mutant animals (Harris and Horvitz, 2011), which we did not investigate. An interesting possibility is that, akin to seam cell division, LIN-29a or LIN-29b may suffice to delay the occurrence of the next molt rather than to prevent it entirely, and in all animals.

Finally, although both isoforms (but not MAB-10) are required for wild-type alae formation, their individual loss causes qualitatively different defects. We speculate that this may reflect an involvement of hyp7, in addition to seam cells, in alae formation, either directly or through an effect on seam cells that remains to be elucidated. The fact that MAB-10 function is not required for wild-type alae formation, but enhances precocious alae formation in *lin-41* mutant animals (Harris and Horvitz, 2011), is then in agreement with the idea that it has a generic function in boosting LIN-29 activity, rather than a specific effect on a subset of LIN-29 regulated targets required for only certain processes. We propose that genome-wide identification of the targets of the *lin-29* isoforms and *mab-10* will allow further testing of this notion in the future. Tissue-specific identification of targets in particular would provide a more detailed understanding of when, where and how different J/A events are triggered. Already, the findings presented in this study have helped to revise our understanding of the fundamental regulatory architecture that temporally controls events occurring during the transition from a juvenile to an adult animal.

## SUPPLEMENTARY FIGURES

**Figure S1.**
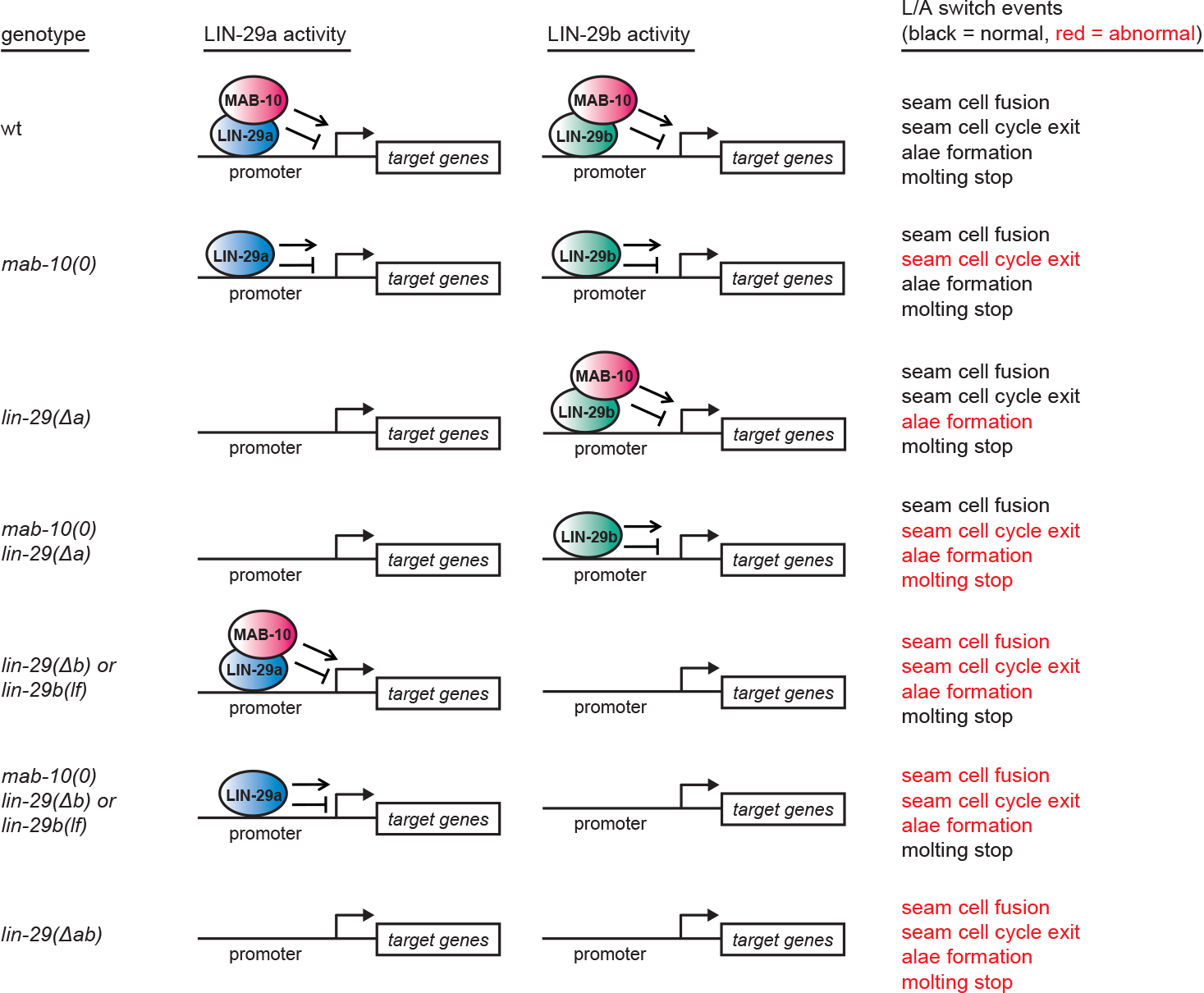
Summary of phenotypes of mutant animals examined in this study. Summary of the J/A transition phenotypes seem for different permutations of *lin-29a*, *lin-29b*, and *mab-10* mutations. Note that extra seam cell divisions in *mab-10(0)* mutant animals occur only in older adults. Some older *mab-10* mutant adults may also undergo extra molts (Harris and Horvitz, 2011), although we failed to observe this.

**Figure S2.**
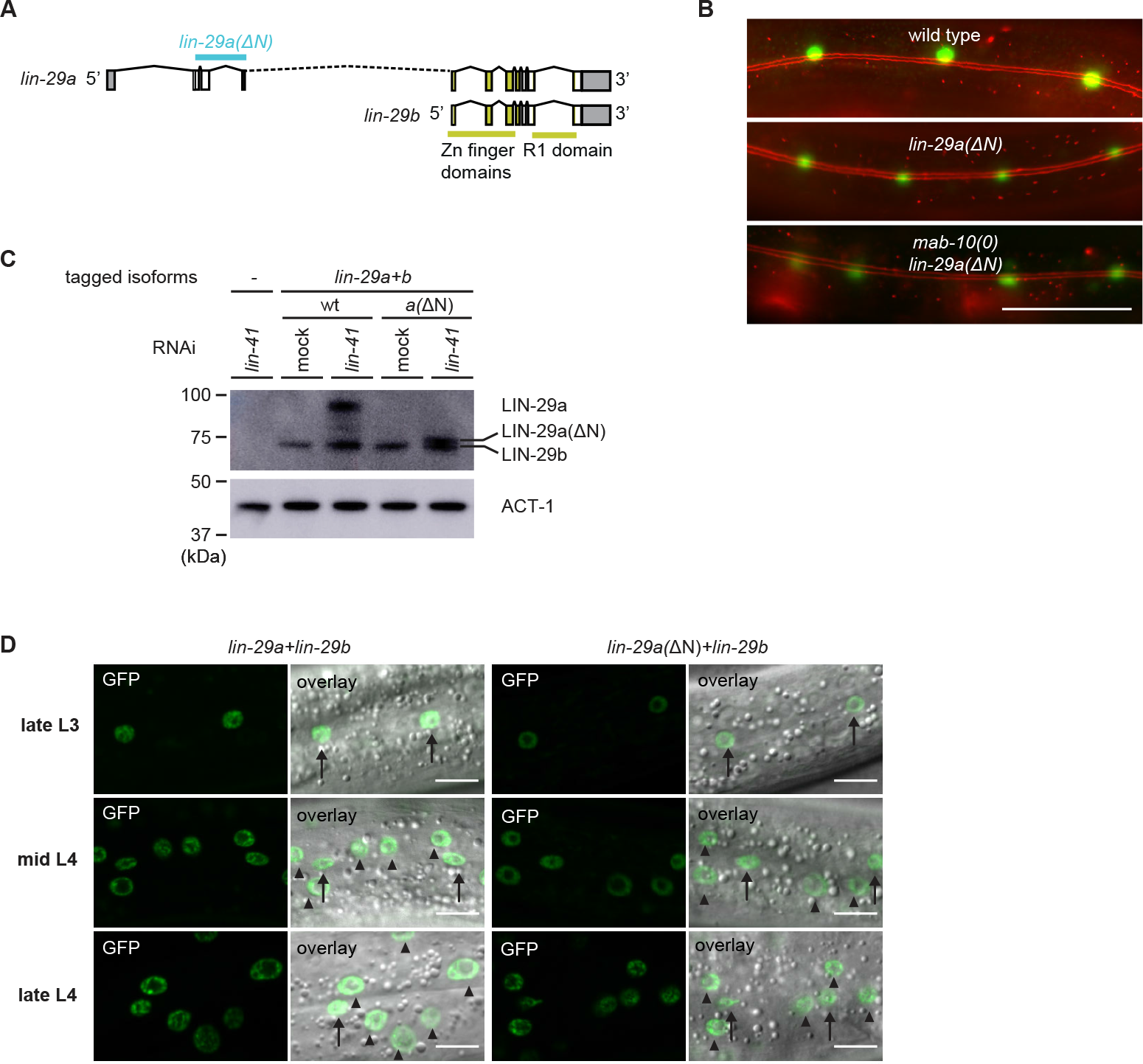
Characterization of *lin-29a(ΔN)* expression and function. A. Schematic representation of the *lin-29a(ΔN) deletion.*
B. Micrographs of late L4-stage animals of indicated genotypes expressing *scm∷gfp* (*green*, marking seam cells) and *ajm-1∷mCherry* (*red*, marking hypodermal cell junctions). In both *lin-29a(ΔN*) and *mab-10(0) lin-29a(ΔN)* animals, fusion occurs normally. Scale bar: 50 μm.
C. Western blot of C-terminally GFP∷3×FLAG-tagged endogenous LIN-29a and LIN-29b proteins in a wild-type and the *lin-29a(ΔN)* background (HW2408), respectively, using anti-FLAG antibody. Animals were grown for 20 h at 25°C to the L3 stage on mock RNAi and *lin-41* RNAi bacteria, respectively. Both LIN-29a and LIN-29a(ΔN) accumulate upon knock-down of *lin-41.* Actin-1 is used as a loading control.
D. Confocal images of endogenously tagged LIN-29 protein isoforms in the wild type and *lin-29a(ΔN)* background (HW2408) in the epidermis of animals at the indicated developmental stages. Animals were staged by examination of gonad development. Arrows indicate seam cell, arrowheads hyp7 nuclei. Scale bars: 10 μm.

**Figure S3.**
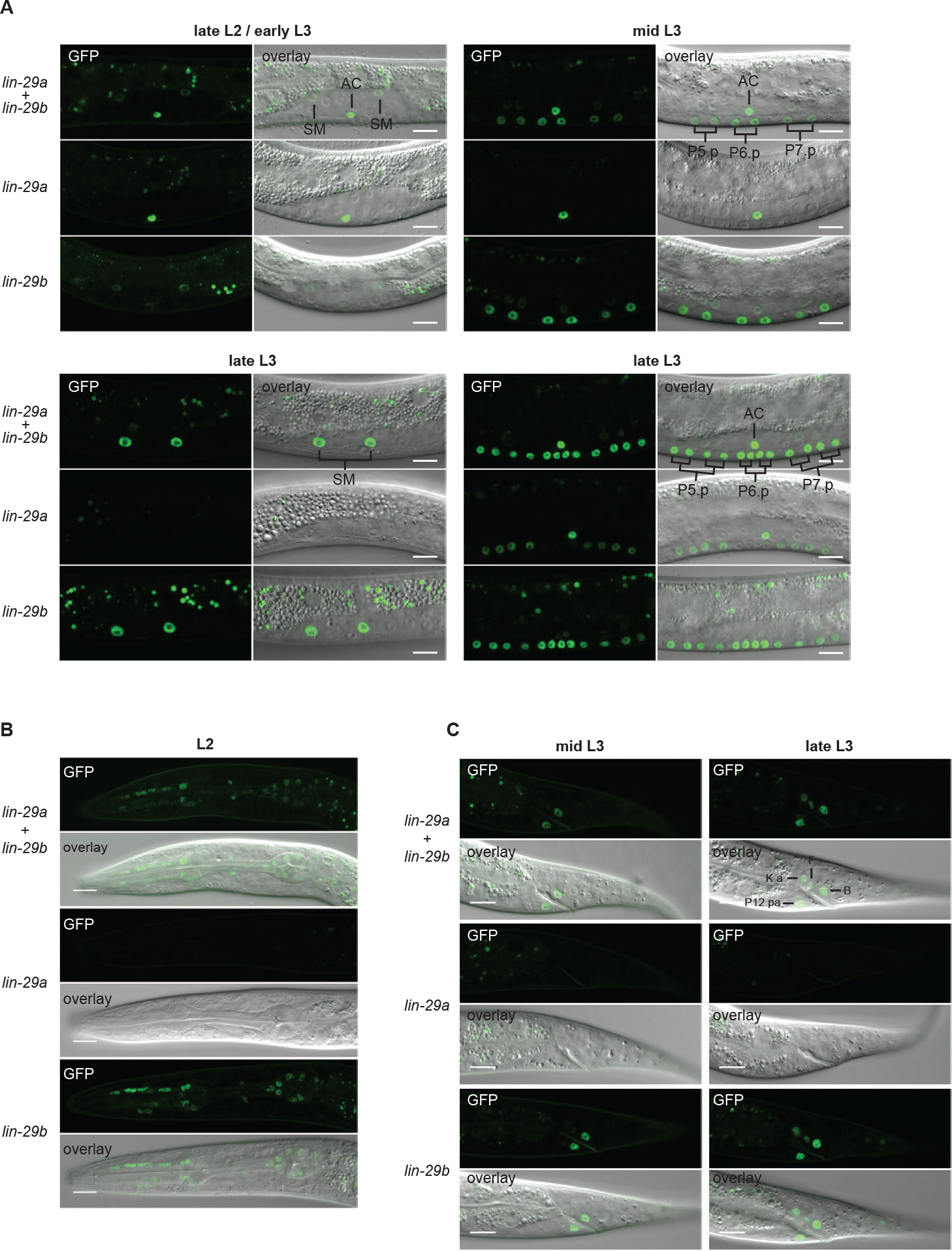
Expression of *lin-29* isoforms. A. Confocal images of endogenously tagged LIN-29 protein isoforms in the region of the vulva and the uterus at the indicated developmental stages. At the L2-to-L3 molt (A), *lin-29a* is expressed in the anchor cell (AC), while *lin-29b* is weakly expressed in the sex myoblasts (SMs). In mid-L3 stage worms, the six daughters of the VPCs P5.p-P7.p express *lin-29b.* At the late L3 stage, *lin-29b* is strongly expressed in the sex myoblast (SM) daughters and all 12 granddaughters of the VPCs P5.p-P7.p, while LIN-29a specifically accumulates in the granddaughters of P5.p and P7.p, but not in those of P6.p. Scale bars: 10 μm.
B. Confocal images of endogenously tagged LIN-29 protein isoforms in the pharynx of L2 stage animals at the indicated developmental stages. *lin-29b* but not *lin-29a* is expressed in the pharynx throughout larval and adult development. Scale bars: 10 μm.
C. Confocal images of endogenously tagged LIN-29 protein isoforms in the tail region at the indicated developmental stages. LIN-29b first accumulates in the two rectal cells B and P12.pa, before accumulating in the four additional rectal cells F, K.a, K’ and U (B, the latter two are not visible in this focal plane). LIN-29a is not detected in these cells. Scale bars: 10 μm.

## SUPPLEMENTARY TABLES

**Table S1:**
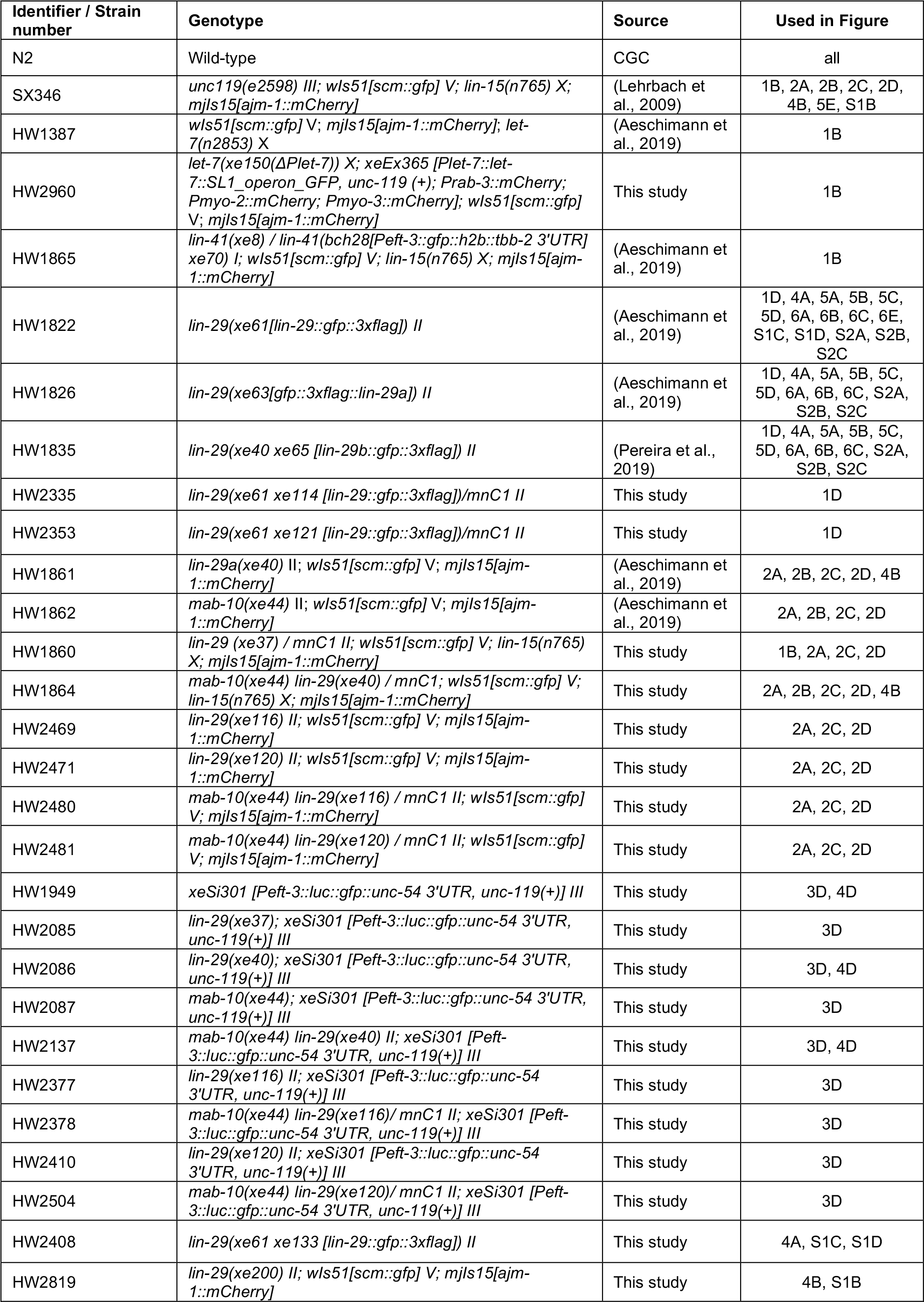
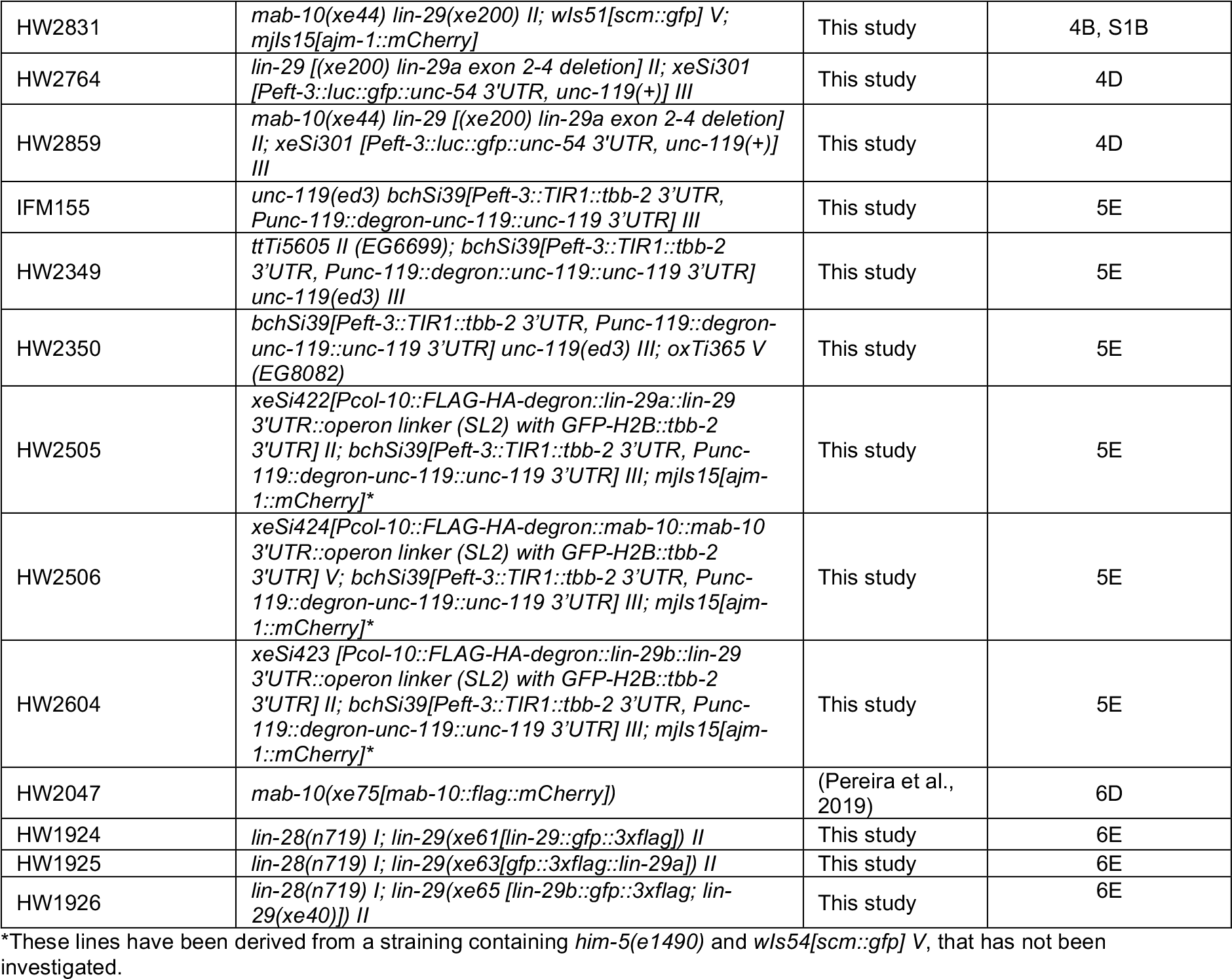
Worm strains used in this study.

**Table S2, S3, S4 and S5: Quantification of phenotypes – related to Figure 1, 2 and 3**

These tables are provided in one combined Excel file.

**Table S6:**
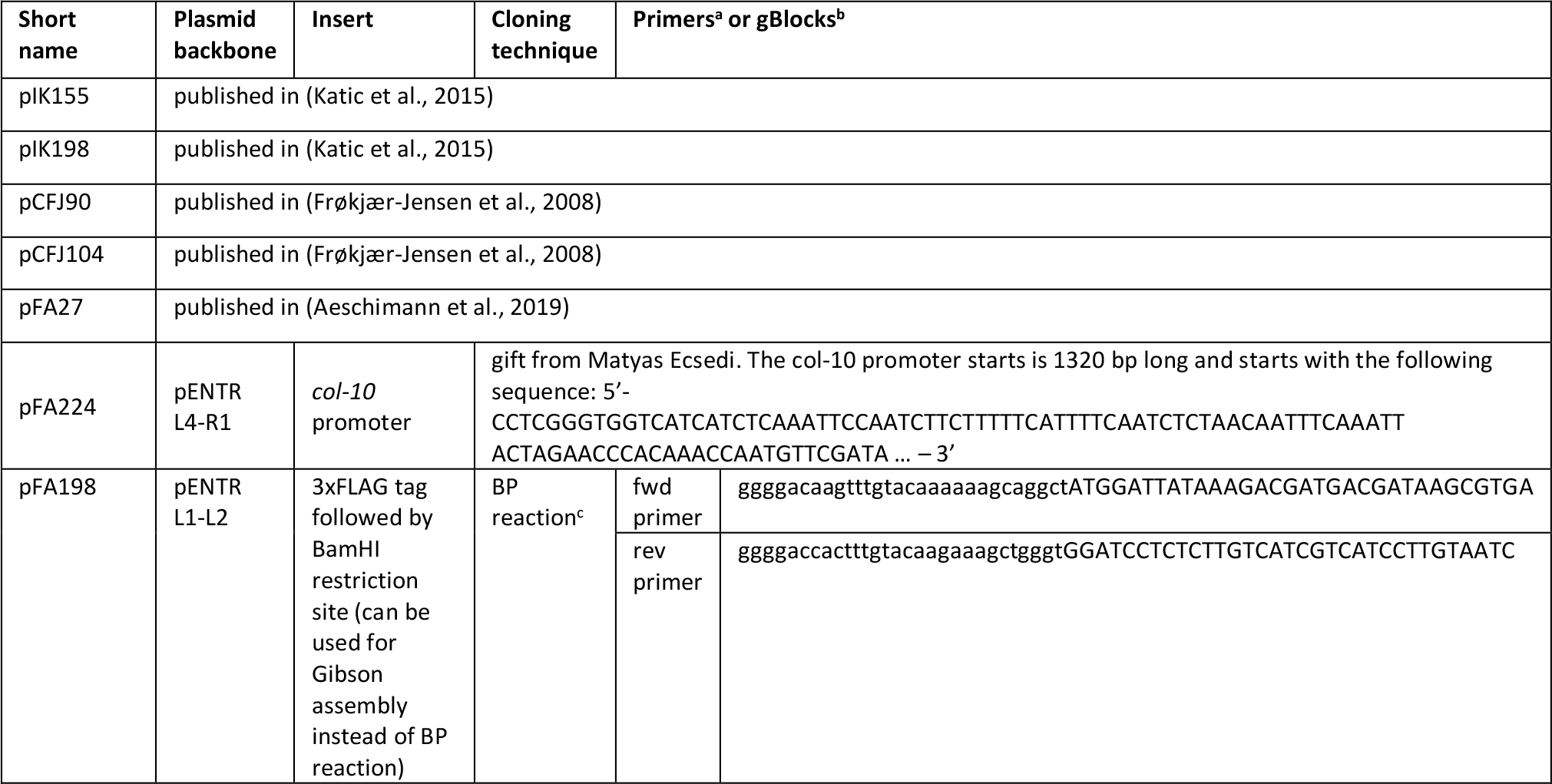
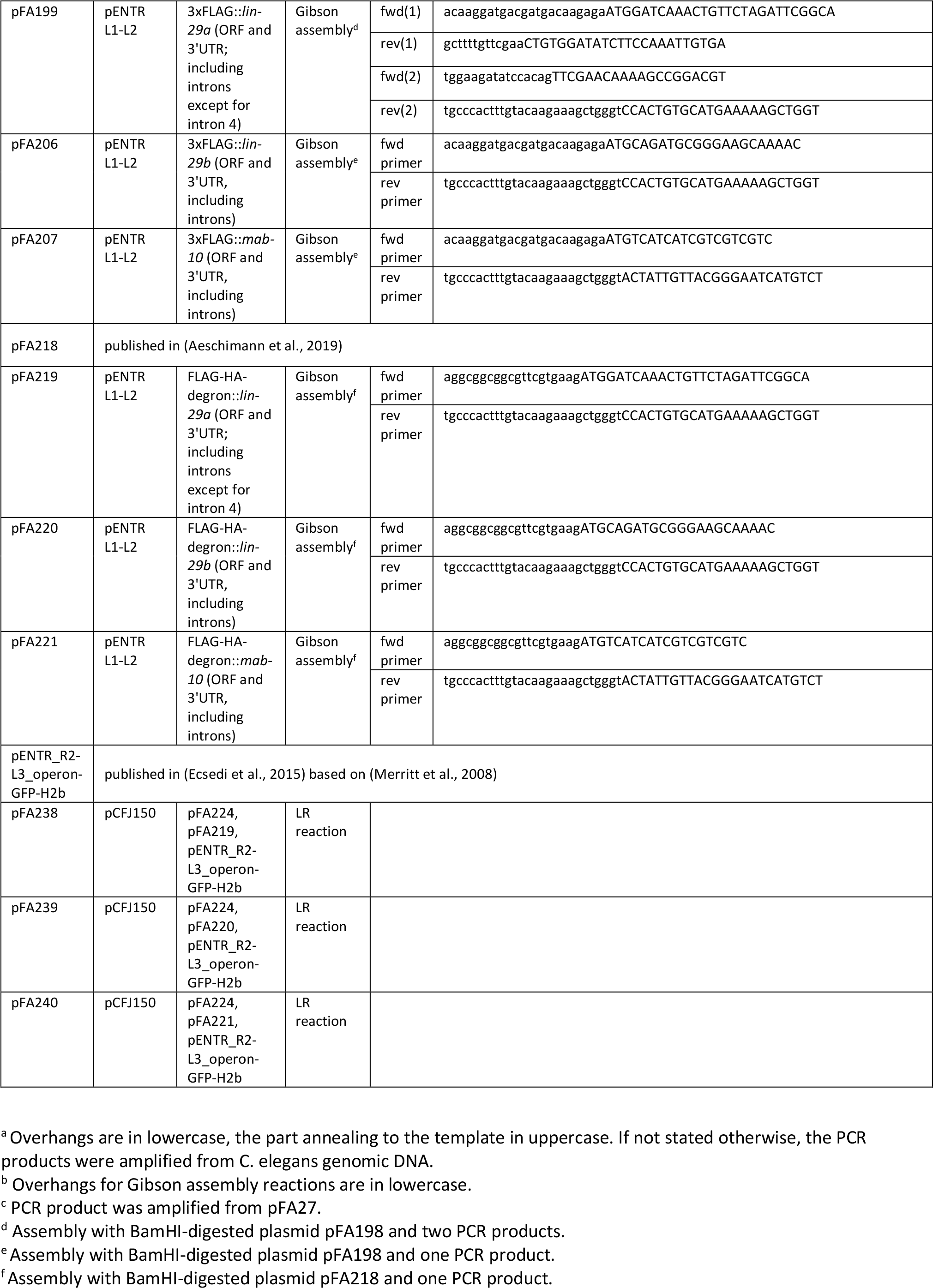
Plasmid used in this study.

## METHODS

### C. elegans

Worm strains used in this study are listed in Table S1. Bristol N2 was used as the wild-type strain. To obtain arrested L1 larval stage worms, embryos were extracted from gravid adults using a bleaching solution (30% (v/v) sodium hypochlorite (5% chlorine) reagent (Thermo Fisher Scientific; 419550010), 750 mM KOH). The embryos were incubated overnight in the absence of food in order to obtain hatched L1 arrsted larvae, at room temperature in M9 buffer (42 mM Na_2_HPO_4_, 22 mM KH_2_PO_4_, 86 mM NaCl, 1 mM MgSO_4_). If not otherwise specified, worms were grown on 2% NGM agar plates with *Escherichia coli* OP50 bacteria (Stiernagle, 2006). For Western blot experiments, worms were grown on enriched peptone plates with *Escherichia coli* NA22 bacteria (Evans, 2006). For RNAi experiments, arrested L1s were plated on RNAi-inducing NGM agar plates with *Escherichia coli* HT115 bacteria containing plasmids targeting the gene of interest (Ahringer, 2006).

### Generation of novel *lin-29b* and *lin-29a* mutant alleles using CRISPR-Cas9

Wild-type worms were injected with a mix containing 50 ng/μl of pIK155, 100 ng/μl of each pIK198 with a cloned sgRNA template, 5 ng/μl pCFJ90 and 5 ng/μl pCFJ104, as previously described (Katic et al., 2015). Single F1 progeny of injected wild-type worms were picked to individual plates and the F2 progeny screened for deletions or point mutations using PCR assays. After analysis by DNA sequencing, the alleles were outcrossed three times to the wild-type strain.

In order to specifically mutate *lin-29b* without affecting expression of *lin-29a*, we generated two different alleles: *lin-29(xe116)* deletes the putative *lin-29b* promoter region to disrupt *lin-29b* transcription, and *lin-29(xe120)* introduces an upstream ATG start codon, resulting in out-of-frame translation on the *lin-29b* mRNA when the ribosome initiates translation from the upstream introduced site.

By injecting two sgRNAs (sgRNA1: acagaattaaggagacaagg, sgRNA2: agctcaattgatctaggttt) into wild-type animals, we obtained the *lin-29(xe116)* allele, a 6724 bp deletion spanning the intron between exon 4 and exon 5, the site of the putative *lin-29b* promoter. The deletion has the following flanking sequences: 5’ atacggtttcagaattaaggagaca – *xe116* deletion – cctagatcaattgagctctaaagat 3’. Injection of the same sgRNAs into *lin-29(xe61[lin-29∷gfp∷3×flag])* generated *lin-29(xe61 xe114)*, which contained the identical mutation in the context of a GFP-3×FLAG-tagged LIN-29.

By injecting two sgRNAs (sgRNA1: gttcgaacaaaagccggacg, sgRNA2: agctgaagcacccccacgtc) into wild-type animals, we obtained the *lin-29(xe120)* allele, by introduction of three point mutations in exon 5. The first mutation introduces an ATG start codon upstream of the normal *lin-29b* ATG start codon. The other two point mutations improve the Kozak context around the introduced ATG codon to increase the efficiency of aberrant translation initiation. All three mutations are silent with respect to the *lin-29a* ORF. The relevant sequence of exon 5 is: TTCGAACAAAAGCC(G=>A)GA(C=>T)GT(G=>C). Injection of the same sgRNAs into *lin-29(xe61[lin-29∷gfp∷3×flag])* generated *lin-29(xe61 xe121)*, which contained the identical mutation in the context of a GFP-3xFLAG-tagged LIN-29.

In order to specifically obtain a *lin-29a* mutant lacking the N-terminal specific domain, we generated the allele *lin-29(xe200)*. By injecting four sgRNAs (sgRNA1: gctggaaccaccactggctc, sgRNA2: gtggcaggagagaattctga, sgRNA3: agccaacttcttcaacgcaa, sgRNA4: gtgaaaacatatgatgtggc), we obtained a 1187 nt deletion spanning exons 2-4 that results in translation of a LIN-29a protein lacking the first 119 amino acids. 23 N-terminal amino acids remain upstream of the LIN-29b N-terminus start codon. The deletion has the following flanking sequences: 5’ ccaacttcttcaacgcaatg - 1187 nt deleted – ttttcacaatttggaagata 3’. Injection of the same sgRNAs into *lin-29(xe61[lin-29∷gfp∷3×flag])* generated *lin-29(xe61 xe133)*, which contained the identical mutation in the context of a GFP-3×FLAG-tagged LIN-29.

### Construction of plasmids for single-copy integrations

All final and intermediate plasmids as well as the primers used in these clonings are listed in Table S2. The genomic regions containing coding exons (from the ATG start codon until the end of the 3’UTR) for *mab-10*, *lin-29a* and *lin-29b* were amplified by PCR using Phusion High-Fidelity DNA Polymerase (NEB, M0530S) from purified genomic wild-type *C. elegans* DNA. The *lin-29a* region was amplified in two fragments, the first spanning exons 2-4 (from ATG in exon 2), the second spanning exons 5-11 (including the 3’UTR). The large intron 4 was not amplified and exons 4 and 5 were fused into one exon when combining the two fragments during Gibson assembly*. lin-29a* 5’UTR was excluded as it is non-coding and confers LIN-41-mediated regulation (Aeschimann et al., 2017).

Using Gibson assembly, the PCR products were cloned into the BamHI site of pFA198. Plasmid pFA198 is a pENTR L1-L2 backbone plasmid that was constructed using Gateway cloning (BP reaction) to insert a 3xFLAG tag followed by a BamHI restriction site for subsequent Gibson assembly. Similarly, a FLAG-HA-degron tag followed by a BamHI restriction site was inserted into a pENTR L1-L2 backbone to obtain pFA218, a template for Gibson assembly reactions (Aeschimann et al., 2019). Plasmids with *mab-10*, *lin-29a* and *lin-29b* genomic regions cloned into pFA198 (i.e., pFA199, pFA206, pFA207) were only used as cloning intermediates in this study. From those plasmids, the *mab-10*, *lin-29a* and *lin-29b* genomic regions were re-amplified by PCR using Phusion High-Fidelity DNA Polymerase and inserted into BamHI-digested pFA218 using Gibson assembly. The resulting plasmids pFA219, pFA220 and pFA221 were then used for further Gateway cloning (LR reactions). In those LR reactions, transgenes were combined with the *col-10* promoter upstream (Table S6) and an operon linker with GFP-H2B∷tbb-2 3'UTR (Merritt et al., 2008) downstream and cloned into the destination vector pCFJ150 (Frøkjær-Jensen et al., 2008). The resulting plasmids pFA238, pFA239 and pFA240 were used for injections.

### Construction of worm strains precociously expressing LIN-29 isoforms or MAB-10

Worm lines with integrated transgenes were obtained by single-copy integration into chromosome II (ttTi5605 locus) or chromosome V (oxTi365), using a protocol for injection with low DNA concentration (Frøkjær-Jensen et al., 2012). In order to perform injections while inducing auxin-mediated degradation of proteins expressed from the introduced transgenes, new Mos1 insertion strains HW2349 and HW2350 were generated. To obtain these strains, the Mos1 insertion strains EG6699 and EG8082 were crossed to worms expressing TIR1 from the *eft-3* promoter (IFM155). Injected animals were grown on NGM plates supplemented with 1 mM auxin (Sigma-Aldrich, I2886). Injection mixes contained pCFJ601 (*Peft-3∷transposase*) at 10 ng/ul, pGH8 (*Prab-3∷mCherry*) at 10 ng/ul, pCFJ90 (*Pmyo-2∷mCherry*) at 2.5 ng/ul, pCFJ104 (*Pmyo-3∷mCherry*) at 5 ng/ul and the respective targeting vector at 50 ng/μl in water. Integrated transgenic lines were further crossed with worms expressing the *ajm-1∷mCherry* marker.

### Seam cell and alae imaging and quantification

Arrested L1 larvae were plated on OP50 bacteria and synchronized worms were grown at 25°C for 36-38 hours (late L4 stage) or 40-42 hours (young adult stage), with the developmental time assessed by staging of individual worms according to gonad length and vulva morphology. Worms were mounted to a 2 % (w/v) agarose pad and immobilized in 10 mM levamisol (Fluca Analytical, 31742). Fluorescent and Differential Interference Contrast (DIC) images were acquired with a Zeiss Axio Observer Z1 microscope using the AxioVision SE64 software and Zen 2 (blue edition). Selections of regions and processing of images was performed with Fiji (Schindelin et al., 2012). Seam cell quantifications were performed by counting all clearly visible fluorescent cells expressing an *scm∷gfp* transgene (Koh and Rothman, 2001) of the upper lateral side in mounted worms. Seam cell fusion quantification was performed by counting seam cells junctions visible through expression of an *ajm-1∷mCherry* transgene (Lehrbach et al., 2009). Alae quantification was performed by observing alae structures with Differential Interference Contrast (DIC) with a 100x objective. For worm lines containing mnC1-balanced animals, *myo-2p*∷GFP-positive (i.e, balancer carrying) animals were excluded from imaging and quantifications. To score *let-7(xe150)* homozygous animals within a population of balanced *let-7(xe150)*; *xeEx365 [Plet-7∷let-7∷SL1_operon_GFP, unc-119 (+); Prab-3∷mCherry; Pmyo-2∷mCherry; Pmyo-3∷mCherry]* animals, all *Pmyo-2∷mCherry* expressing (i.e., balancer carrying) animals were excluded from the analysis. To score *lin-41(xe8)* homozygous animals within a population of balanced *lin-41(xe8)*/*lin-41(bch28 xe70)* animals, all *eft-3p*∷*gfp∷h2b* expressing (i.e., balancer carrying) animals were excluded from the analysis.

### Luciferase assay

Luciferase assays were performed as described (Meeuse et al., 2019). Briefly, embryos were extracted from gravid adults using a bleaching solution. Single embryos were transferred into a well of a white, flat-bottom, 384-well plate (Berthold Technologies, 32505) by pipetting. Animals were left to develop in 90 uL S-Basal medium containing *E. coli* OP50 (OD_600_ = 0.9) and 100 μM Firefly D-Luciferin (p.j.k., 102111). Plates were sealed with Breathe Easier sealing membrane (Diversified Biotech, BERM-2000). Luminescence was measured using a luminometer (Berthold Technologies, Centro XS3 LB 960) every 10 minutes for 0.5 sec for 90/100 hours in a temperature controlled incubator set to 20 degrees. Luminescence data was analyzed using an automated algorithm to detect the hatch and the molts in MATLAB, as described before (Meeuse et al., 2019).

### Confocal imaging

For detection of endogenously tagged LIN-29∷GFP, GFP∷LIN-29a and LIN-29b∷GFP (HW1826, HW1882, HW1835) by confocal microscopy, animals were grown at 25°C on OP50 bacteria. For confocal imaging of endogenously tagged LIN-29∷GFP, GFP∷LIN-29a and LIN-29b∷GFP (HW1826, HW1882, HW1835) and MAB-10∷mCherry (HW2047) on RNAi conditions, synchronized arrested L1 stage larvae were grown for 20 hours at 25 °C on RNAi-inducing plates with HT115 bacteria. The bacteria either contained the L4440 parental RNAi vector without insert (denoted “mock RNAi”) or with an insert targeting *lin-41* (Fraser et al., 2000) or *hbl-1* (Kamath and Ahringer, 2003). Worms were imaged on a Zeiss LSM 700 confocal microscope driven by Zen 2012 Software after mounting them on a 2% (w/v) agarose pad with a drop of 10 mM levamisol solution. Differential Interference Contrast (DIC) and fluorescent images were acquired with a 40×/1.3 oil immersion objective (1024×1024 pixels, pixel size 156nm).

For confocal imaging of endogenously tagged LIN-29∷GFP, GFP∷LIN-29a and LIN-29b∷GFP in the wild-type and *lin-28(n719)* mutant background (HW1826, HW1882, HW1835, HW1924, HW1925, HW1926) worms were grown at 25°C on OP50 bacteria for 20 and 22 hours respectively (to accommodate a modest delay in development of *lin-28(n719)* mutants). For confocal imaging of endogenously tagged LIN-29∷GFP in the wild-type and *lin-29a*(*ΔN)* background (HW1826, HW2408), worms were grown at 25°C on OP50 bacteria. In both cases, worms were imaged on an Axio Imager M2 (upright microscope) + Yokogawa CSU W1 Dual camera T2 spinning disk confocal scanning unit driven by Visiview 3.1.0.3 software after mounting them on a 2% (w/v) agarose pad with a drop of 10 mM levamisol solution. Differential Interference Contrast (DIC) and fluorescent images were acquired with a 40×/1.3 oil immersion objective (2048×2048 pixels, 16-bits). Using the Fiji software (Schindelin et al., 2012), images were processed after selecting representative regions. Worms of the same worm line were imaged and processed with identical settings.

### Testing for precocious seam cells fusion in worms precociously expressing LIN-29 isoforms or MAB-10

All strains were grown on OP50-seeded NGM plates supplemented with 1 mM auxin. The strain expressing MAB-10 from the *col-10* promoter (HW2506) was grown on *mab-10* RNAi plates supplemented with auxin (100 ug/ml carbenicillin, 1 mM IPTG, 1 mM auxin). Gravid animals were bleached and eggs were incubated at room temperature in M9 for hatching overnight. To allow accumulation of LIN-29a, LIN-29b or MAB-10, the animals were not exposed to auxin anymore from this point onward. Synchronized L1 animals were plated and analyzed on NGM plates with OP50 at 25°C. Seam cell fusion was scored by observation of the *ajm-1∷mCherry* marker at 15h after plating synchronized L1 animals.

### Western blotting

Animals were grown for 20 hours at 25°C on RNAi-inducing plates, as described above for confocal imaging, or on NA22 plates. Lysates were made by boiling (5 minutes, 95°C) and sonication in SDS lysis buffer (63 mM Tris-HCl (pH 6.8), 5 mM DTT, 2% SDS, 5% sucrose) and cleared by centrifugation, before separating proteins by SDS-PAGE (loading: 50 μg protein extract per well) and transferring them to PVDF membranes by semi-dry blotting. The following antibodies were used: Monoclonal mouse anti-FLAG M2-Peroxidase (HRP) (Sigma-Aldrich; A8592, dilution: 1:1,000). Monoclonal mouse anti-Actin clone C4 (Millipore; MAB1501, dilution 1:10,000). Detection was performed with a horseradish peroxidase-conjugated secondary antibody (NXA931), ECL Western Blotting Detection Reagents and an ImageQuant LAS 4000 chemiluminescence imager (all from GE Healthcare).

## QUANTIFICATION AND STATISTICAL ANALYSIS

All values of n indicate numbers of animals. No statistical tests were performed.

## AUTHOR CONTRIBUTION

C.A. conceived part of the project; designed, performed, and analyzed confocal microscopy, luciferase assay and western blot experiments; created transgenic worm lines; and wrote the manuscript. F.A. conceived the project; designed, performed, and analyzed confocal microscopy, phenotypic analysis and western blot experiments; created transgenic worm lines; and edited the manuscript. A.N. performed and analyzed the precious expression experiment; and created transgenic worm lines. H.G. conceived the project; designed and analyzed experiments; acquired funding; and wrote the manuscript.

## Acknowledgements

We thank Milou Meeuse for the gift of HW1949 and help with the luciferase assay, Jana Kracmarova for the gift of *let-7(xe150)*, Iskra Katic for the gift of IFM155, and Foivos Gypas for computational analysis. Some strains were provided by the Caenorhabditis Genetics Center, which is funded by the NIH Office of Research Infrastructure Programs (P40 OD010440). We thank Iskra Katic, Jana Kracmarova for a critical reading of the manuscript. We are particularly grateful to Benjamin Towbin for extensive comments, suggestions and discussions. We thank Lan Xu and Iskra Katic for technical support. This project was supported through funding from the Swiss National Science Foundation (#31003A_163447) and Friedrich Miescher Institute for Biomedical Research core funding through the Novartis Research Foundation (to H.G.).

## REFERENCES

Abete-Luzi, P., and Eisenmann, D.M. (2018). Regulation of C. elegans L4 cuticle collagen genes by the heterochronic protein LIN-29. Genesis 56.

Abrahante, J.E., Daul, A.L., Li, M., Volk, M.L., Tennessen, J.M., Miller, E.A., and Rougvie, A.E. (2003). The Caenorhabditis elegans hunchback-like gene lin-57/hbl-1 controls developmental time and is regulated by microRNAs. Dev. Cell 4, 625–637.

Abreu, A.P., Dauber, A., Macedo, D.B., Noel, S.D., Brito, V.N., Gill, J.C., Cukier, P., Thompson, I.R., Navarro, V.M., Gagliardi, P.C., et al. (2013). Central precocious puberty caused by mutations in the imprinted gene MKRN3. N. Engl. J. Med. 368, 2467–2475.

Aeschimann, F., Kumari, P., Bartake, H., Gaidatzis, D., Xu, L., Ciosk, R., and Großhans, H. (2017). LIN41 Post-transcriptionally Silences mRNAs by Two Distinct and Position-Dependent Mechanisms. Mol. Cell 65, 476–489.e4.

Aeschimann, F., Neagu, A., Rausch, M., and Großhans, H. (2019). Let-7 coordinates the transition to adulthood through a single primary and four secondary targets. Life Sci. Alliance 2, 1–18.

Ahringer, J. (2006). Reverse Genetics. WormBook, Ed. C. Elegans Res. Community, Wormb.

Ambros, V. (1989). A hierarchy of regulatory genes controls a larva-to-adult developmental switch in C. elegans. Cell 57, 49–57.

Ambros, V. (2011). MicroRNAs and developmental timing. Curr. Opin. Genet. Dev. 21, 511–517.

Ambros, V., and Horvitz, H.R. (1984). Heterochronic mutants of the nematode Caenorhabditis elegans. Science (80-.). 226, 409–416.

Bettinger, J.C., Lee, K., and Rougvie, A.E. (1996). Stage-specific accumulation of the terminal differentiation factor LIN-29 during Caenorhabditis elegans development. Development 122, 2517–2527.

Bettinger, J.C., Euling, S., and Rougvie, A.E. (1997). The terminal differentiation factor LIN-29 is required for proper vulval morphogenesis and egg laying in Caenorhabditis elegans. Development 124, 4333–4342.

Chalfie, M., Horvitz, H.R., and Sulston, J.E. (1981). Mutations that lead to reiterations in the cell lineages of C.elegans. Cell 24, 59–69.

Corre, C., Shinoda, G., Zhu, H., Cousminer, D.L., Crossman, C., Bellissimo, C., Goldenberg, A., Daley, G.Q., and Palmert, M.R. (2016). Sex-specific regulation of weight and puberty by the Lin28/let-7 axis. J. Endocrinol. 228, 179–191.

Cox, G.N., and Hirsh, D. (1985). Stage-specific patterns of collagen gene expression during development of Caenorhabditis elegans. Mol. Cell. Biol. 5, 363–372.

Ding, X.C., and Großhans, H. (2009). Repression of C. elegans microRNA targets at the initiation level of translation requires GW182 proteins. EMBO J. 28, 213–222.

Ecsedi, M., Rausch, M., and Großhans, H. (2015). The let-7 microRNA directs vulval development through a single target. Dev. Cell 32, 335–344.

Evans, T. (2006). Transformation and microinjection. WormBooked, Ed. C. Elegans Res. Community, Wormb.

Faunes, F., and Larraín, J. (2016). Conservation in the involvement of heterochronic genes and hormones during developmental transitions. Dev. Biol. 416, 3–17.

Fraser, A.G., Kamath, R.S., Zipperlen, P., Martinez-Campos, M., Sohrmann, M., and Ahringer, J. (2000). Functional genomic analysis of C. elegans chromosome I by systematic RNA interference. Nature 408, 325–330.

Frøkjær-Jensen, C., Wayne Davis, M., Hopkins, C.E., Newman, B.J., Thummel, J.M., Olesen, S.P., Grunnet, M., and Jorgensen, E.M. (2008). Single-copy insertion of transgenes in Caenorhabditis elegans. Nat. Genet. 40, 1375–1383.

Frøkjær-Jensen, C., Davis, M.W., Ailion, M., and Jorgensen, E.M. (2012). Improved Mos1-mediated transgenesis in C. elegans. Nat. Methods 9, 117–118.

Großhans, H., Johnson, T., Reinert, K.L., Gerstein, M., and Slack, F.J. (2005). The temporal patterning microRNA let-7 regulates several transcription factors at the larval to adult transition in C. elegans. Dev. Cell 8, 321–330.

Harris, D.T., and Horvitz, R.H. (2011). MAB-10/NAB acts with LIN-29/EGR to regulate terminal differentiation and the transition from larva to adult in C. elegans. Development 138, 4051–4062.

Hunter, S.E., Finnegan, E.F., Zisoulis, D.G., Lovci, M.T., Melnik-Martinez, K. V., Yeo, G.W., and Pasquinelli, A.E. (2013). Functional Genomic Analysis of the let-7 Regulatory Network in Caenorhabditis elegans. PLoS Genet. 9.

Kamath, R.S., and Ahringer, J. (2003). Genome-wide RNAi screening in Caenorhabditis elegans. Methods 30, 313–321.

Katic, I., Xu, L., and Ciosk, R. (2015). CRISPR/Cas9 genome editing in Caenorhabditis elegans: Evaluation of templates for homology-mediated repair and knock-ins by homology-independent DNA repair. G3 Genes, Genomes, Genet. 5, 1649–1656.

Koh, K., and Rothman, J.H. (2001). ELT-5 and ELT-6 are required continuously to regulate epidermal seam cell differentiation and cell fusion in C. elegans. Development 128, 2867–2880.

Lee, P.A., and Houk, C.P. (2007). Puberty and its disorders.

Lehrbach, N.J., Armisen, J., Lightfoot, H.L., Murfitt, K.J., Bugaut, A., Balasubramanian, S., and Miska, E.A. (2009). LIN-28 and the poly(U) polymerase PUP-2 regulate let-7 microRNA processing in Caenorhabditis elegans. Nat. Struct. Mol. Biol. 16, 1016–1020.

Lin, S.Y., Johnson, S.M., Abraham, M., Vella, M.C., Pasquinelli, A., Gamberi, C., Gottlieb, E., and Slack, F.J. (2003). The C. elegans hunchback homolog, hbl-1, controls temporal patterning and is a probable MicroRNA target. Dev. Cell 4, 639–650.

Meeuse, M.W.M., Hauser, Y.P., Hendriks, G., Eglinger, J., Bogaarts, G., Tsiairis, C., and Großhans, H. (2019). State transitions of a developmental oscillator. Biorxiv. Https://Doi.Org/10.1101/755421.

Merritt, C., Rasoloson, D., Ko, D., and Seydoux, G. (2008). 3ʹ UTRs Are the Primary Regulators of Gene Expression in the C. elegans Germline. Curr. Biol. 18, 1476–1482.

Moss, E.G., and Romer-Seibert, J. (2014). Cell-intrinsic timing in animal development. Wiley Interdiscip. Rev. Dev. Biol. 3, 365–377.

Myster, D.L., and Duronio, R.J. (2000). Cell cycle: To differentiate or not to differentiate? Curr. Biol. 10, 302–304.

Olmedo, M., Geibel, M., Artal-Sanz, M., and Merrow, M. (2015). A high-throughput method for the analysis of larval developmental phenotypes in Caenorhabditis elegans. Genetics 201, 443–448.

Ong, K.K., Elks, C.E., Li, S., Zhao, J.H., Luan, J., Andersen, L.B., Bingham, S.A., Brage, S., Smith, G.D., Ekelund, U., et al. (2009). Genetic variation in LIN28B is associated with the timing of puberty. Nat. Genet. 41, 729–733.

Pereira, L., Aeschimann, F., Wang, C., Lawson, H., Serrano-Saiz, E., Portman, D.S., Großhans, H., and Hobert, O. (2019). Timing mechanism of sexually dimorphic nervous system differentiation. Elife 8, 1–31.

Perry, J.R.B., Stolk, L., Franceschini, N., Lunetta, K.L., Zhai, G., McArdle, P.F., Smith, A. V., Aspelund, T., Bandinelli, S., Boerwinkle, E., et al. (2009). Meta-analysis of genome-wide association data identifies two loci influencing age at menarche. Nat. Genet. 41, 648–650.

Reinhart, B.J., Slack, F.J., Basson, M., Pasquienelll, A.E., Bettlnger, J.C., Rougvle, A.E., Horvitz, H.R., and Ruvkun, G. (2000). The 21-nucleotide let-7 RNA regulates developmental timing in Caenorhabditis elegans. Nature 403, 901–906.

Rougvie, A.E., and Ambros, V. (1995). The heterochronic gene lin-29 encodes a zinc finger protein that controls a terminal differentiation event in Caenorhabditis elegans. Development 121, 2491–2500.

Rougvie, A.E., and Moss, E.G. (2013). Developmental transitions in C. elegans larval stages. Curr. Top. Dev. Biol. 105, 153–180.

Schindelin, J., Arganda-Carreras, I., Frise, E., Kaynig, V., Longair, M., Pietzsch, T., Preibisch, S., Rueden, C., Saalfeld, S., Schmid, B., et al. (2012). Fiji: An open-source platform for biological-image analysis. Nat. Methods 9, 676–682.

Singh, J.E., and Sulston, R.N. (1978). Some observation on moulting in Caenorhabditis elegans. Nematologica 63–71.

Slack, F.J., Basson, M., Liu, Z., Ambros, V., Horvitz, H.R., and Ruvkun, G. (2000). The lin-41 RBCC gene acts in the C. elegans heterochronic pathway between the let-7 regulatory RNA and the LIN-29 transcription factor. Mol. Cell 5, 659–669.

Stiernagle, T. (2006). Maintenance of C.elegans. WormBook, Ed. C. Elegans Res. Community, Wormb.

Sulem, P., Gudbjartsson, D.F., Rafnar, T., Holm, H., Olafsdottir, E.J., Olafsdottir, G.H., Jonsson, T., Alexandersen, P., Feenstra, B., Boyd, H.A., et al. (2009). Genome-wide association study identifies sequence variants on 6q21 associated with age at menarche. Nat. Genet. 41, 734–738.

Sulston, J.E., and Horvitz, H.R. (1977). Post-embryonic cell lineages of the nematode, *Caenorhabditis elegans*. Dev Biol 56, 110–156.

Vadla, B., Kemper, K., Alaimo, J., Heine, C., and Moss, E.G. (2012). Lin-28 controls the succession of cell fate choices via two distinct activities. PLoS Genet. 8.

Van Wynsberghe, P.M., Kai, Z.S., Massirer, K.B., Burton, V.H., Yeo, G.W., and Pasquinelli, A.E. (2011). LIN-28 co-transcriptionally binds primary let-7 to regulate miRNA maturation in Caenorhabditis elegans. Nat. Struct. Mol. Biol. 18, 302–308.

Zhang, L., Ward, J.D., Cheng, Z., and Dernburg, A.F. (2015). The auxin-inducible degradation (AID) system enables versatile conditional protein depletion in C. elegans. Dev. 142, 4374–4384.

Zhu, H., Shah, S., Shyh-Chang, N., Shinoda, G., Einhorn, W.S., Viswanathan, S.R., Takeuchi, A., Grasemann, C., Rinn, J.L., Lopez, M.F., et al. (2010). Lin28a transgenic mice manifest size and puberty phenotypes identified in human genetic association studies. Nat. Genet. 42, 626–630.

